# Dose-Efficient Cryo-Electron Microscopy for Thick Samples using Tilt-Corrected Scanning Transmission Electron Microscopy, Demonstrated on Cells and Single Particles

**DOI:** 10.1101/2024.04.22.590491

**Authors:** Yue Yu, Katherine A. Spoth, Michael Colletta, Kayla X. Nguyen, Steven E. Zeltmann, Xiyue S. Zhang, Mohammadreza Paraan, Mykhailo Kopylov, Charlie Dubbeldam, Daniel Serwas, Hannah Siems, David A. Muller, Lena F. Kourkoutis

## Abstract

Cryo-EM is a powerful tool in structural biology, providing insights through techniques like single-particle analysis (SPA) and cryogenic electron tomography (cryo-ET). In thick specimens, challenges arise as an exponentially larger fraction of the transmitted electrons lose energy from inelastic scattering and can no longer be properly focused as a result of chromatic aberrations in the post-specimen optics. Rather than filtering out the inelastic scattering at the price of reducing potential signal, as is done in energy-filtered transmission electron microscopy (EFTEM), we show how a dose-efficient and unfiltered image can be rapidly obtained using tilt-corrected bright-field scanning-TEM (tcBF-STEM) data collected on a pixelated detector. Enhanced contrast and a 3-5x improvement in collection efficiency are observed for 2D images of intact bacterial cells and large organelles using tcBF-STEM compared to EFTEM for thicknesses beyond 500 nm. As a proof of concept for the technique’s performance in structural determination, we present an SPA map at subnanometer resolution for a highly symmetric virus-like particle (VLP) with 789 particles. These findings suggest applications for tcBF-STEM in cryo-EM of thicker cellular volumes where current approaches struggle.

## Main

Cryogenic electron microscopy (cryo-EM) provides powerful insights into the study of biological systems by revealing molecular structures in their close-to-native environments^1–3^. Single particle analysis (SPA) has enabled structural determination of purified macromolecular complexes up to atomic resolution^4,5^. Cryogenic electron tomography (cryo-ET) with subtomogram averaging (STA) has been developed to resolve macromolecular structures in biological contexts including within slices of whole cells^6,7^. Compared to SPA, fewer structures have been resolved at high resolution by cryo-ET with STA with one of the main limitations being the increased specimen thickness for cellular structures compared to the preparations for the purified molecules. This increased sample thickness leads to an exponential decrease in the elastically-scattered signal, especially at high sample tilts^8^ or lower beam voltages^9^. In the conventional transmission electron microscopy (TEM) geometry, the imaging optics are placed after the sample, and chromatic blur in the post-specimen optics leads to a strong defocusing of the inelastically scattered electrons. Energy-filtered TEM (EFTEM) removes this blur caused by inelastic scattering but in doing so reduces the collected signal and dose-efficiency compared to an ideal microscope^10,11^. Chromatic aberration correction could in principle correct some of this inelastic blur over a limited energy range and it is an ongoing topic of active research to improve the energy range, stability and resolution^12,13^.

It has also long been recognized that in the scanning transmission electron microscopy (STEM) geometry, where the electron beam is focused to a small spot and then rastered across the specimen, that post-specimen chromatic aberrations should not compromise the probe size. This is because in STEM the probe-forming optics are placed before the sample, and before any inelastic scattering can occur, thus STEM imaging should be less susceptible to specimen-induced chromatic blurring (instead the chromatic blur in the post-specimen optics degrades the angular coherence of the diffraction pattern). Consequently, the possibility of studying µm-thick biological samples with STEM tomography has been explored both experimentally and theoretically, utilizing coherent, incoherent signals and a combination of both^14–18^.

Recent advances in the design of STEM detectors^19–22^ have enabled rapid 4D-STEM data acquisition, where almost all of the scattered electrons are collected as 2D images of convergent beam electron diffraction (CBED) patterns, and recorded over a 2D grid of probe positions, as sketched in Fig.1a. 4D-STEM has simplified the implementation of other STEM phase imaging techniques such as (integrated) differential phase contrast (iDPC-)STEM^23^ and electron ptychography^24–26^. Efforts have been made to optimize these techniques for applications in structural biology studies. iDPC-STEM has generated the first SPA map of macromolecules embedded in vitrified ice by a STEM technique at near-atomic resolution^23^. Initial attempts at low-dose ptychography have been performed on purified virus-like-particles (VLPs) at nanometer resolution with a limited number of particles^24,25^. More recent demonstrations have shown that SPA of thin sections with ptychography can resolve protein structures at a sub-nanometer level^26^, including a 5.8Å SPA map of apoferritin reconstructed from ∼11,000 particles. This performance is still worse than EFTEM and TEM when beam-induced motion is corrected, and suggests the resolution limit is not the instrument optics, but likely related to uncorrected sample motion under the beam. To date, both the iDPC and ptychography studies have focused on relatively thin samples that were optimized for SPA applications.

Here we describe how, in STEM geometry, a new dose-efficient phase-contrast imaging technique—tilt-corrected bright-field (tcBF-) STEM—could prove useful for imaging thick samples, while still providing comparable spatial resolution for thin samples. With this technique, we were able to resolve features in thick samples (roughly 500-800 nm thick) that were not visually discernible with EFTEM under comparable conditions in intact bacterial cells and large organelles. Additionally, with single particle approach, we present a ∼7 Å nominal resolution 3D map for a highly symmetric virus-like particle (VLP) from 789 particles as proof of feasibility for structural determination with tcBF-STEM. Our earlier work on tcBF can be found in a series of short conference abstracts^27–31^ but a detailed writeup of the method had been delayed by the illness and untimely passing of our colleague Lena Kourkoutis, and here we provide a more in-depth description. This technique is computationally much faster than iterative ptychography, so could be used for live monitoring while collecting 4D-STEM data. We note this technique is starting to find applications in the development of low-dose ptychography for cryo-EM applications and materials science studies^26,32^.

The starting point for tcBF-STEM is the collection of a 4D-STEM data set (Fig.1a), similar to what might be recorded for an out-of-focus ptychographic reconstruction^33^. For tcBF-STEM, each pixel within the bright-field (BF) disk functions as a coherent BF detector subtending a sufficiently small collection angle. From the theorem of reciprocity^34^, the STEM image produced from the detector pixel on the optical axis is equivalent to a conventional BF TEM image, and those STEM images produced by off-axis detector pixels are equivalent to BF TEM images formed with tilted illumination (Fig. 1b). These equivalent beam tilts give rise to image shifts that depend on the aberration function^35,36^, and are particularly simple when the dominant aberration is defocus. (There are some important differences for inelastic scattering^37^ that we discuss below and more details are given in the online methods section, where we follow the image analysis framework laid out by Rose^37^.) Such an image shift is demonstrated with two images obtained with two off-axis detector pixels (Fig. 1c-d). The shifts are measured and corrected on a (detector) pixel-by-pixel basis. Fig. 1e and 1g illustrate the resolved shift map overlaid on the averaged CBED pattern. Each individual image, after shift correcting, is then combined to create the final tcBF-STEM image (Fig. 1h). Compared to the BF images formed by single detector pixel (Fig. 1d), the tcBF-STEM image has a significantly improved SNR because almost all the signal-relevant signals are utilized. Furthermore, compared to the image formed by directly integrating over the full BF disk (Fig. 1f), tcBF preserves phase contrast. When reconstructing a tcBF-STEM image, a simultaneous measurement of the probe aberration function can be obtained. In fact, one of the early applications of a shift analysis of 4D-STEM datasets was for aberration measurement^38^, by analogy with the TEM beam tilt methods^35^. In tcBF-STEM, like in conventional BF-TEM, defocus is deliberately introduced to enhance contrast. Consequently, in Fig.1 e and g, the magnitudes are linearly proportional to the defocus and the off-axis angles, and are oriented outwards. The linearity of the shift with angle also makes it possible to measure the depth of objects by the resulting parallax effect^22^.

**Figure 1.**
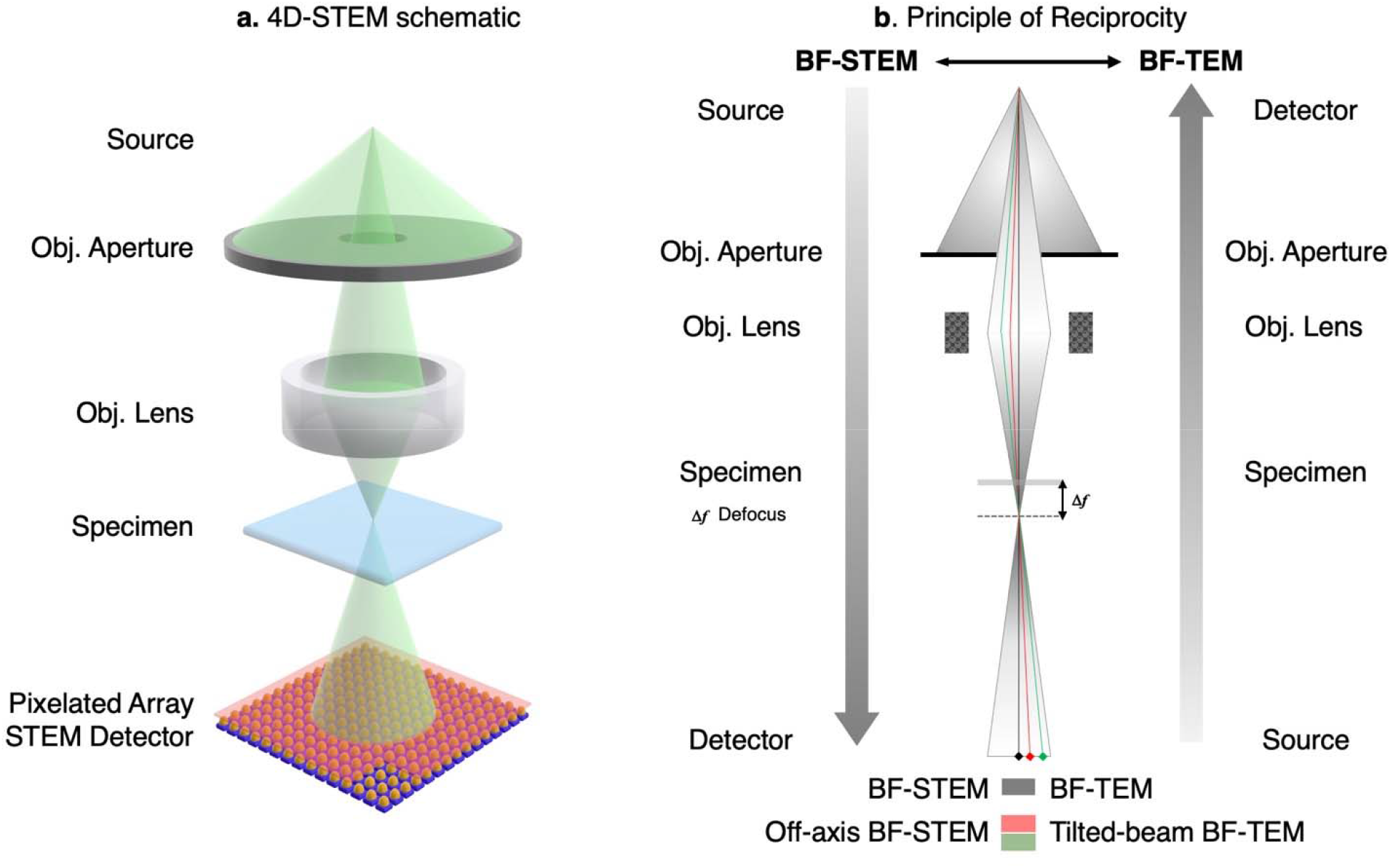
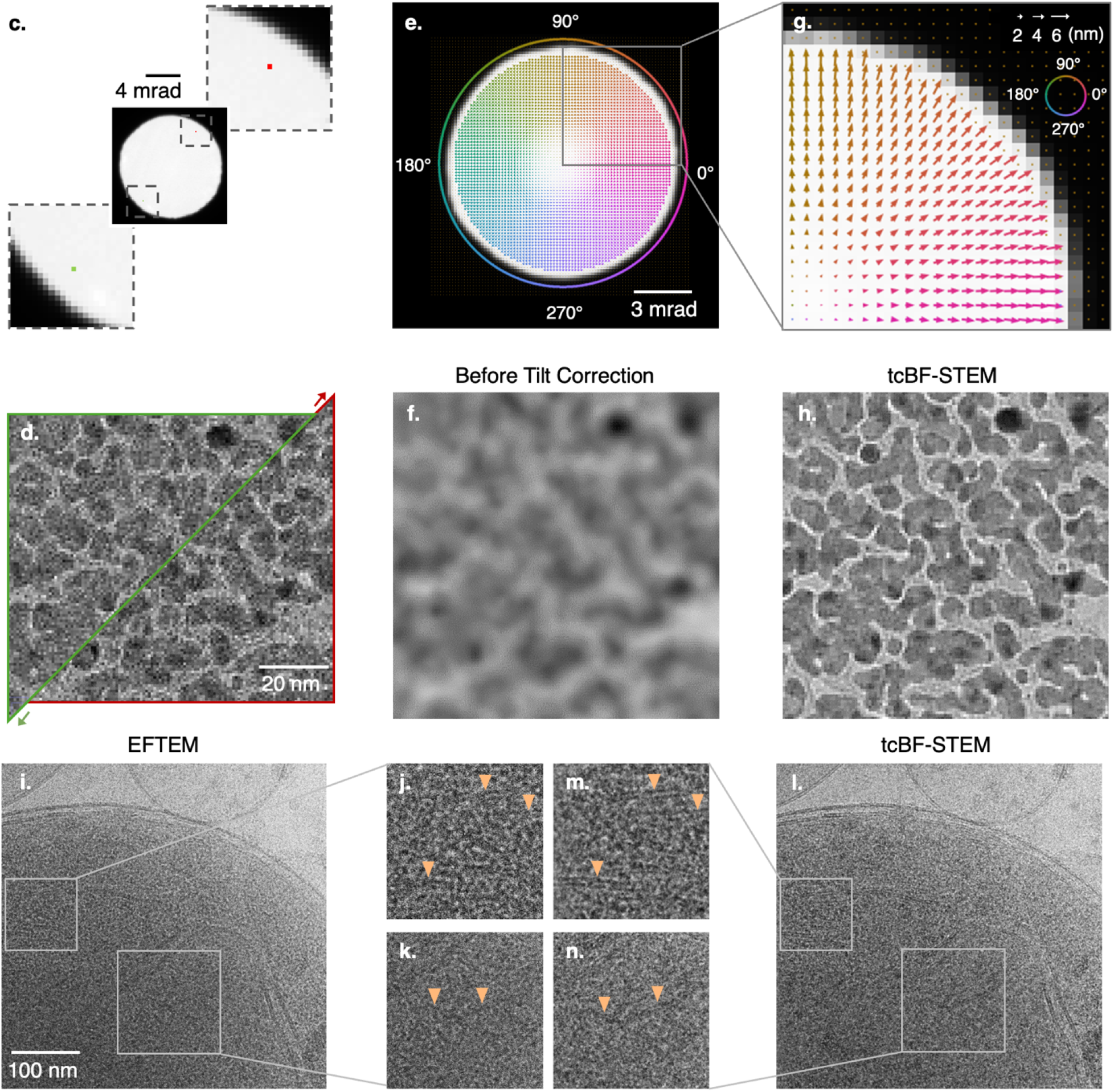
Direct phase-contrast imaging with 4D STEM: tilt-corrected bright-field (tcBF) -STEM employs a pixelated STEM detector to collect the entire convergent beam electron diffraction (CBED) pattern (a). Each detector pixel within the bright-field (BF) disk is a coherent BF STEM detector though located off the optical axis. By reciprocity (b), off-axis BF-STEM (top down) is equivalent to tilted-illumination TEM (bottom up). Similar to BF TEM, defocus is applied to introduce phase contrast. For a standard gold-on-carbon sample and a defocused probe, integrating the signals collected by two off-axis detector pixels (red and green) in (c) produces two images with relative shifts between them (d). For every detector pixel, the image shifts determined through cross-correlation with the on-axis detector pixel are shown by the arrows overlaid on the averaged CBED pattern in (e) with a zoom-in and binned view in (g). The arrows are color-coded corresponding to the shift directions. Integrating the full forward-scattered bright-field (BF) signals without correcting for the angle-dependent shifts results in blurring (f) due to the defocus. A tcBF-STEM image (h) is generated by summing the images after shift correction. In a tcBF-STEM image, the signal-to-noise ratio (SNR) is increased compared to (d) and the blurring due to defocus (f) is corrected. To compare the performance of tcBF-STEM with energy-filtered TEM (EFTEM) on thick samples, (i) and (l) are the images acquired in the same area in a mitochondrion. The dose measured over vacuum is 14 e^-^/Å^2^, and the acceleration voltage is 300 kV for both acquisitions. More information can be found in table I. The membrane bilayer was similarly resolved in the thin part of the sample for both methods. However, in thicker regions (j) and (m), tcBF still shows the membrane bilayers clearly (indicated by the orange arrowheads), while this feature is less visible with EFTEM. In the even thicker region (k and n), tcBF can still resolve some parts of the inner membranes whereas with EFTEM these features are less discernible. A thickness estimate map, obtained from the fraction of electrons remaining in the energy-filtered image, is given in Fig. S1, with thicknesses ranging from 470 nm to 620 nm

We are now also in a position to understand the challenges for dose-efficient STEM with a single-pixel detector, and why tcBF-STEM overcomes that. By reciprocity, the conventional TEM geometry would be reproduced in STEM with a single small-pixel detector on the optic axis. The smaller the angular range of the detector, the more coherent the signal – in TEM mode, this would be equivalent to the illumination angle. But in STEM mode, such a small detector collects only a tiny fraction of the incident beam – a 0.1 mrad wide collector, and a 10 mrad probe convergence angle would have a collection efficiency of 1 part in 10,000, whereas a TEM with a 0.1 mrad illumination convergence, and a 10 mrad post-specimen objective aperture would have almost perfect collection efficiency. To improve the collection efficiency in STEM, we could increase the collection angle of the detector but this will eliminate the phase contrast signal (a much weaker amplitude contrast in an incoherent image will still be present – e.g. chapter 3 of reference 39). This is because the phase-contrast signal is only measurable when there is a phase shift on the lens, but the phase shift from aberrations generates an image shift that is different for each angle. In other words, simply summing over a wide range of angles leads to a blurred image. If the dominant aberrations are defocus and coma, the images recorded on the off-axis detector pixels have similar contrast transfer functions to the on-axis pixel^39^ (except towards the edge of the aperture – full treatment in online methods), so the tilt-correct summation of tcBF corrects for these shifts, allowing a coherent image to be retained, and uses almost all of the incident beam -i.e. a similar dose efficiency to TEM. The presence of the aperture complicates the analysis compared to aperture-free TEM, but the end result for tcBF is a similar-looking contrast transfer function (CTF) that has an information limit at double the aperture size (see results and online methods). This is the same information limit cutoff for iDPC and bright-field ptychography, although the shapes of the CTF are very different. As we will discuss in the results section, iDPC is less efficient than tcBF at transferring low-frequency information, although it is simpler to interpret.

Moreover, tcBF-STEM has an advantage over EFTEM for thick samples. The post-specimen lenses for EFTEM are the image formation lenses so chromatic aberrations in the post-specimen lenses degrade the image resolution. However, for tcBF-STEM, the post-specimen lenses simply transfer an image of the diffraction pattern so chromatic aberrations result in a small loss of angular resolution -i.e. a small increase in the effective detector pixel size and hence a reduction in coherence. In a thick sample, most electrons undergo both elastic and inelastic scattering, but the elastic contrast is preserved when scattered to the inelastic channels^37,40^. This is largely because the most-likely inelastic scattering events are very delocalized compared to the elastic scattering, leading to weak and low-frequency modulations of the real space signal, and only a small (∼0.03-0.1 mrad) blurring of the angular distributions.

For a qualitative comparison of EFTEM and tcBF-STEM in thick specimens, we imaged the same area in a mitochondrion in succession with the two techniques (Fig.1i-n) using the same incident dose of 14 e^-^/Å^2^, and the same acceleration voltage of 300 kV. In the thinnest part of the sample at the organelle’s edge, the membrane bilayers are similarly resolved with both methods. However, in thicker regions (Fig.1 j and m), tcBF-STEM clearly shows the bilayers (orange arrow) of the mitochondrial inner membranes whereas with EFTEM these features are less visible. In the thickest portion of the image, tcBF can still resolve some parts of the inner membranes (Fig.1 k) whereas in the EFTEM image (Fig.1 n) these features are hardly discernible. Using the unfiltered and 10-eV EFTEM images and the inelastic mean free path (MFP) for vitrified ice of ∼310 nm^41^ at 300 kV, the sample’s thickness can be estimated (see online methods). At the mitochondrion’s edge, the sample thickness is approximately 500 to 520 nm thick, while the regions shown in Fig. 1j-m is about 570 to 600 nm, and Fig. 1k-n corresponds to around 600 to 620 nm (thickness map in Fig S1). In the thickest parts (k-n), the EFTEM signal has dropped to ∼14% of the incident dose, but the tcBF still retains 50% of the incident dose, i.e. almost 3.6x more signal remaining for tcBF. Similar trends are also observed in multiple samples shown later in Fig.3, where the differences in collection efficiencies are compared quantitatively, with the relative efficiency of tcBF-STEM over EFTEM growing exponentially as the sample thickness increases.

## Results

### Fast Data Acquisition with tcBF-STEM Upsampling and the CTF of tcBF-STEM

In tcBF-STEM the number of pixels in the reconstructed image can be made much larger than the number of diffraction patterns recorded, as at a finite defocus each diffraction pattern contains information about an extended region of the sample. This trade-off between real space and reciprocal space sampling helps speed up the data collection as the multi-pixel detectors used for tcBF tend to be slower to readout than single-pixel sensors. For instance, if each diffraction pattern records information about 8*8 subsampled regions, the data collection rate is sped up 64-fold, so a 10 kHz detector frame rate becomes a 640 kHz image-pixel rate. The recovery of information beyond the limits of the real-space probe sampling uses the information collected in the shadow images in the diffraction plane. The information retrival is achieved by a real-space upsampling through sub-(scan)-pixel image shifting. To understand the implementation of this upsampling technique, we start with a demonstration on a standard gold-on-carbon sample. This same approach is then also applied in the data reconstruction workflow for all the examples shown in the paper.

Figure 2 shows a tcBF-STEM dataset acquired with 256*256 scan positions, spaced 8Å apart. With the chosen defocus value and scan step size (see the online method section), we expect over 90% information overlap collected in the reciprocal space. This information surplus is used to upsample a tcBF image. Details on upsampling can be found in online methods and Fig.S2 to S4. Figure 2a and 2b demonstrate the process of upsampling with part of the detector pixels. Fig. 2d is the final upsampled result with sub-scan-pixel features resolved compared to the original image (c). In Fig. 2e, Thon rings and the 2.3-Å spacing of Au are recovered through upsampling. An FFT radial average profile (g) is shown to confirm that upsampling restores the information beyond the scan Nyquist frequency without altering the information within the frequency range. The upsampling procedure achieves information transfer up to 7 times the real-space Nyquist sampling limit, effectively speeding up data acquisition by a factor of 49.

**Figure 2.**
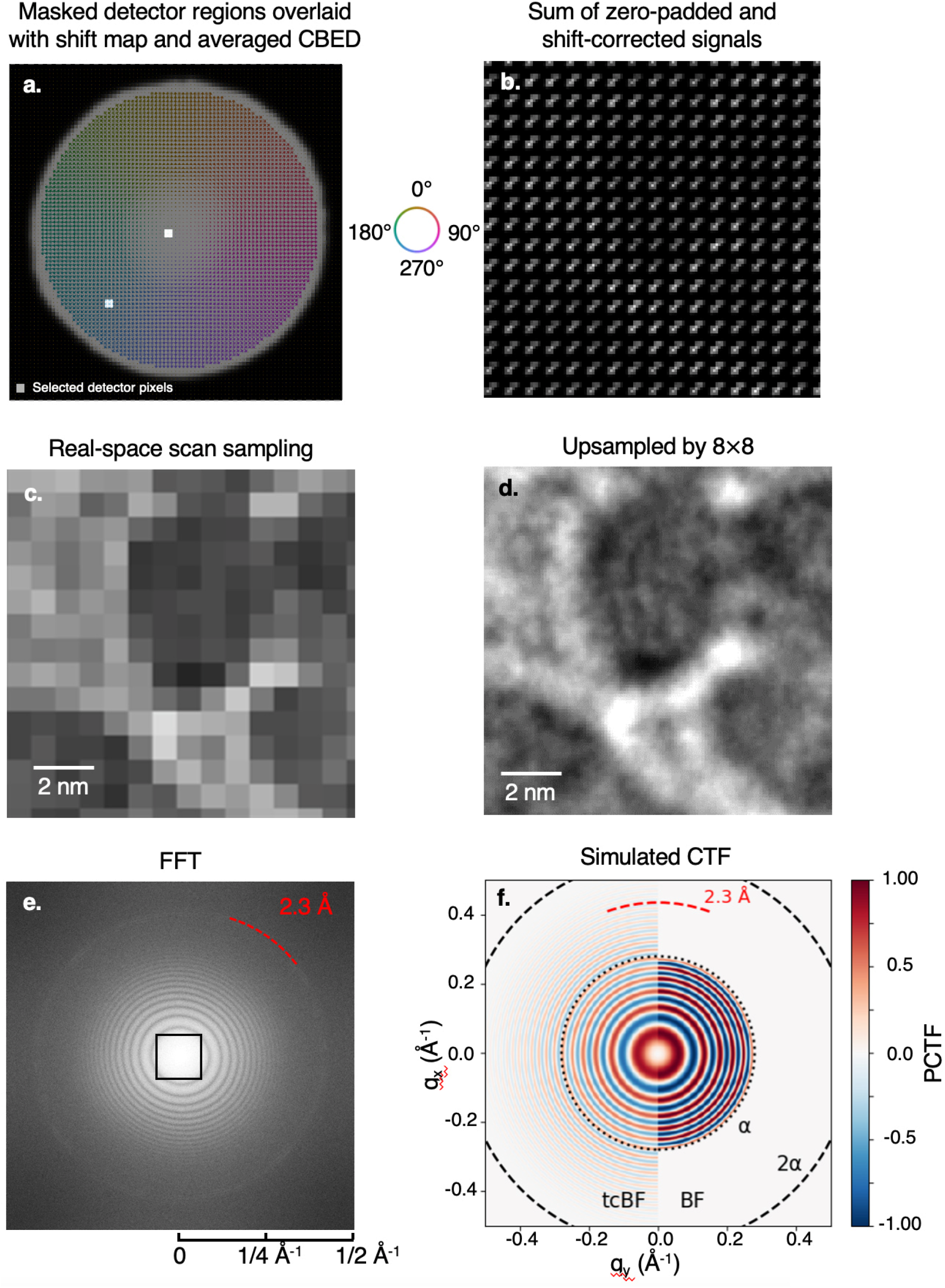
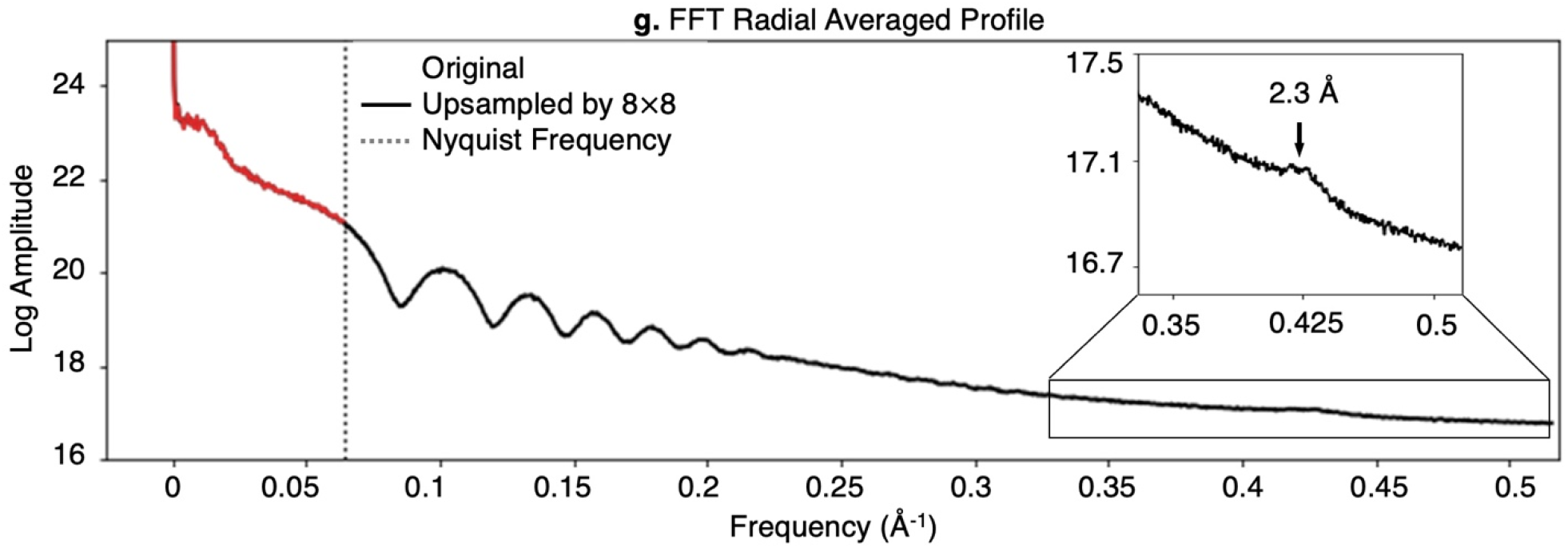
Up-sampling by 8 speeds up data acquisition by 64-fold. Up-sampling for tcBF-STEM of a gold-on-carbon combined test sample is accomplished by exploiting the image shifts from different detector pixels as a result of defocus (and higher order aberrations). (a) The colored arrows show the shift measured for the scanned images synthesized at each detector pixel inside the bright field disk. Scanned images formed by the two white pixels on the detector shown in (a) will be shifted from each other. Correcting for these shifts and accumulating signals collected from the selected detector regions fills in different regions of the scanned image (b) at a spacing finer than the recorded probe positions, demonstrating the first step of a complete up-sampling. A tcBF-STEM image (c) is collected with a defocused probe at an 8-Å scan step size. In the up-sampled tcBF-STEM image (d), additional sub-scan-pixel features are resolved compared to the original image (c). (e) Experimental power spectrum from the full image of the test sample showing Thon rings and the 2.3-Å ring of gold lattice spacing beyond the scan Nyquist frequency (1/16 Å^-1^, black box) are recovered by up-sampling. An FFT radial average profile (g) shows that up-sampling recovers information beyond the scan Nyquist frequency without altering the signal within the electron-optical information limit. (f) The calculated phase contrast transfer function (PCTF) for a tcBF image after shift correction shows twice the information limit compared to the BF image formed using only the axial detector pixel, as a result of exploiting off-axis information. The simulation uses 5.5 mrad convergence semi-angle probe-forming aperture (α), 300 kV acceleration voltage and 700 nm defocus.

The number of pixels in the image is separate from the optical resolution limit. Notably, with the α=5.5-mrad convergence semi-angle, the information transfer limit at a cutoff of 1α corresponds to 3.6 Å and 1.8 Å at 2α. The 2.3-Å spacing we observed exceeds the 1α cutoff and but is just within the 2α limit. As discussed in the previous section, tcBF has a similar-looking PCTF (Fig.2f) to BF TEM but with an information limit at double the aperture size. This is because the information limit is set by the highest spatial frequency that can be transferred, i.e. the maximum possible momentum transfer. For an axial detector, this would be from an incident wavevector on the radius of the probe-forming aperture to the axis. An off-axis detector that is displaced in the opposite direction to the incident wavevector allows for a maximum momentum transfer that spans the diameter of the aperture, doubling the information limit compared to the axial case.

The calculated phase contrast transfer function (PCTF) shown in Fig. 2f shows the tcBF image has twice the information limit compared to the BF image formed using only the axial detector pixel, as a result of exploiting this off-axis information. Details can be found in the online method section and Fig. S5-S7, including a discussion of practical limits.

### Comparison of cryo-tcBF-STEM, conventional TEM, and EFTEM for imaging thick samples

Indeed, we believe tcBF has an advantage for thick specimens. To compare the performance of tcBF-STEM with EFTEM, the most-widely-adopted imaging technique in cryo-EM, we performed successive imaging with the two techniques on various thick specimens, including intact bacterium cells and large cellular organelles. The incident dose is chosen to be the same for each comparison, slit width for EFTEM is 10 eV and the acceleration voltage is 300 kV. As the samples are much thicker than the depth of field, quantitative metrics on the full projection convey less information than they would for thin sections, or individual molecules at different depths (which can be determined by the parallax shift in tcBF). Instead we present comparative cases with different acquisition orders, and different defocus choices for the two techniques.

Even though no high-resolution information is compared, the image acquired first still introduces radiation damage and conformation change prior to the second. Therefore, we present scenarios where EFTEM images were acquired first, as well as scenarios where tcBF-STEM images were acquired first. Additionally, CTF modulation can affect the qualitive comparison. However, achieving the exact same defocus for the two techniques can be challenging because the samples are thick (∼ 550 nm to 700 nm, table) and not flat, and switching between TEM and STEM modes is a significant change in optical alignments. As a result, we present a series of cases where EFTEM images are measured to have defoci larger, equivalent to or smaller than those of tcBF, alongside a scenario where both techniques are targeted at the same nominal defocus.

In Fig.1 (i-n), we showed a comparison on a mitochondrion where tcBF performs better at resolving inner membrane especially towards the thicker part of the organelle. This tcBF image was acquired first with less measured defocus (1.9μm, Fig.3 table I) than EFTEM (3.9μm). EFTEM defoci are measured with CTFFIND4^42^ and tcBF defoci are measured from the image shifts. In this case, membrane contrast in the images is compared across the two techniques but the orientation of the membrane relative to the beam can affect its contrast so differences might be a result of warping of the sample. Another possibility is that the inherent range of tilts in tcBF illumination (up to ∼7 mrad) improves contrast for a larger range of membrane orientations. But in general, we observe improved contrast in thick specimen regions for tcBF where other features instead of membranes were compared. Fig.3a-b are EFTEM and tcBF images of an intact E.coli cell. With tcBF, features within the cell’s interior (Fig.3d with arrows) are effectively resolved, while in EFTEM the same features (possibly condensates or surface contamination) are discernible but less prominent. For this comparison, EFTEM has a lesser value of measured defocus (3.7 μm) than STEM (4.2 μm). In (e) and (f), images with tcBF and EFTEM of a vesicle were acquired with very close defoci (2.8μm). Again, the features in the thick region are clearer compared to EFTEM. We attribute the improved contrast with tcBF to a more efficient use of electrons. In table (I) we compare the ratio of preserved electrons with the two methods. For the samples measured here, tcBF is observed to collect more than 3 times the number of electrons than EFTEM for the same incident dose. For another comparison on E. coli at a low dose of 0.5 e-/Å^2^ (i-l), tcBF is capable of resolving features that are otherwise indiscernible with EFTEM. Table I provides a summary of information on specimen, acquisition orders, doses, pixel sizes, measured or nominal defocus, thickness estimate using the EFTEM image, and a comparison of dose efficiency.

**Table I.** Summary for the information on specimen, doses, pixel sizes, measured or nominal defocus, thickness estimate using the inelastic MFP, and a comparison of the dose efficiency of EFTEM and tcBF.

|  | Specimen | Acquired first | Dose (e<sup>-</sup>/Å<sup>2</sup>) | Pixel size (Å) |  | Defocus (nm) |  | Average thickness estimate* (nm) | Fraction of incident electrons in the image |  |
| --- | --- | --- | --- | --- | --- | --- | --- | --- | --- | --- |
|  |  |  |  | EFTEM | tcBF | EFTEM | tcBF |  | EFTEM | tcBF |
| Fig.1i-n | mitochondrion | STEM | 14 | 2.37 | 3.59<br>(up-sample 8) | 3978.4 | 1943.6 | 547 | 0.171 | 0.533 |
| Fig.3a-d | E.coli | EFTEM | 14 | 2.37 | 3.59<br>(up-sample 8) | 3743.8 | 4262.9 | 673 | 0.114 | 0.403 |
| Fig.3e-h | vesicle | EFTEM | 14 | 2.37 | 3.59<br>(up-sample 8) | 2849.2 | 2825.0 | 586 | 0.151 | 0.522 |
| Fig.3i-l | E.coli | STEM | 0.5 | 12.96<br>(bin by 4) | 10.8<br>(up-sample 4) | 4000<br>(nominal**) | 4000<br>(nominal) | -- | -- | -- |

All data were acquired with 300 kV acceleration voltage at cryogenic temperature
\* Average thickness is estimated with inelastic scattering MFP using the ratio of electrons in the 10-eV EFTEM image and the incident electrons over vacuum.
\*\* Nominal defocus is the calibrated instrument defocus after focusing using a nearby area

Overall, a common trend is that tcBF is more likely to retain higher SNR features in thick regions of samples compared to EFTEM. Figure 4a shows the measured fraction of electrons collected for tcBF and EFTEM, where tcBF collects a factor of 3-3.5x more signal, an advantage that grows with thickness. We expect both signals to decay exponentially with thickness (t), i.e.exp(-t *λ*_in_) for EFTEM and exp (-t *λ*_el_) Fitting to the tcBF data in table I, we find the Elastic MFP is *λ*_*el*_ =830 ±50 nm of the online methods (Figure 4a). Some sense of the relative dose efficiency of the two approaches is given by the ratio of these two exponential decays, i.e.exp(t/*λ*_eff_)= exp(-t/*λ*_el_) exp(-t/*λ*_in_) where −1/ *λ*_*eff*_=1/ *λ*_*el*_−1/ *λ*_*in*,_ so*λ*_*eff*_ ≈ 500 nm. This gives the factor of 3 advantage for tcBF at 550 nm, and it grows to 5x at ∼800 nm. Beyond a thickness of one elastic MFP, much of the phase contrast signal will be lost to multiple elastic scattering and leaving mostly amplitude contrast. This identifies an effective dose advantage window for tcBF over EFTEM for thicknesses beyond ∼400 nm.

For low spatial frequencies, the collection efficiencies for tcBF and EFTEM can be compared directly because of their similar contrast transfer functions (Figure 4b). STEM methods that also collect the entire bright field disk such as DPC and iDPC can also be compared after accounting for their differences in information transfer as a function of spatial frequency. This is captured by the detective quantum efficiency (DQE) of the imaging system (online methods equations A15-17). Figure 4c shows that tcBF is more efficient at low spatial frequencies. The iDPC CTF and DQE peak at zero defocus and degrade with increasing defocus^43^, unlike EFTEM and tcBF.

To understand the sample damage as a function of dose, we consecutively acquire tcBF-STEM images with doses ranging from 1.5 e^−^ /Å^2^ to 210 e^−^ /Å^2^ (Fig.S8 a-d). After a cumulative exposure of 280 e^−^ /Å^2^, no obvious sign of bubbling is observed, consistent with previous STEM studies^16^, while visible bubbling effects start to form after a total exposure over 150 e^−^ /Å^2^ for conventional TEM^16^. This does not mean no damage has occurred, but rather damage products have not migrated over long length scales. This suggests that for cryo-ET, the STEM operation mode might offer a higher total dose tolerance, but it also depends on the desired resolution of information from a tomogram.

### Single particle analysis 3D reconstruction with cryo-tcBF-STEM imaging

To quantitatively assess the current performance of tcBF-STEM for molecular structure analysis, we performed SPA of bacteriophage PP7 coat protein, achieving a nominal resolution of ∼7 Å at 0.143 FSC cutoff using a generic cryoSPARC SPA workflow^44^. For this analysis 789 particles are extracted from 19 tcBF-STEM micrographs. Fig. 5a displays a cropped representative micrograph with 2D class average of the particles in the inset. Per-micrograph CTF estimation was performed by CTFFIND4^42^ without local refinement due to the limited number of particles. Approximately 200 particles were manually picked to generate the template for template picking, and 789 particles were selected. The selected classes were then used for ab initio model generation, as a starting model for the homogeneous refinement with icosahedral symmetry applied. Fig. 5b presents the cryo-EM density map sharpened with Guinier B factor of 351 Å^2^ based on the Guinier plot analysis of the 3D reconstruction. A zoom-in view of the EM density with X-ray crystal structure of the particle docked inside (Protein Data Bank code 1DWN^45^) is shown in (c). Fourier shell correlation (FSC) indicates a nominal resolution of 7.03 Å with cryoSPARC dynamic mask and 9.6 Å with no mask.

VLP PP7 possesses an icosahedral symmetry with triangulation number T=3, theoretical molecular weight of 2MDa, containing a high number of repeated units per particle which allows efficient structural averaging with a smaller number of particles. ∼7-Å resolution demonstrates the feasibility of using tcBF-STEM for structural analysis at a resolution that can resolve some secondary structures such as alpha helices. We also have preliminary experimental results suggesting no resolution limit improvements with the current ptychography algorithms with a similar number of particles. On the other hand, state-of-art EFTEM imaging under similar doses and the same accelerating voltage can achieve 3.5 Å nominal resolution with 900 particles (Fig. S10). We also attempted iDPC on the same specimen with a 1000kHz segmented detector (Fig. S11) and observed lower-quality phase contrast. We suspect this was mainly due to a poor experimental focus determination as we found iDPC to be very sensitive to focus settings. The supporting Quantifoil material for this sample is gold and the thickness is 50 nm. Low-dose constraints restricted focusing to be only on the supporting foil instead of the sample and the 50-nm-thick supporting foil can introduce a focus offset.

## Discussion

We demonstrate tcBF-STEM imaging on purified single particle VLPs and vitrified cellular specimens. A comparative analysis with EFTEM highlights the higher dose efficiency of tcBF-STEM, particularly for thick specimens. The performance of tcBF-STEM on thick specimens in two scenarios was further demonstrated, under low-dose imaging and under cumulative exposures, suggesting potential advantages of tcBF for cryo-ET applications. As a proof-of-concept for using this technique for structural determination, tcBF SPA with VLPs shows a ∼7 Å nominal resolution 3D map using a generic processing workflow for conventional TEM with ∼ 800 particles.

A key advantage of tcBF is in imaging thick specimens, as it is relatively insensitive to specimen-introduced energy losses. tcBF-STEM stands out as a STEM technique due to its high-dose efficiency, as it takes advantage of nearly all the forward-scattering electrons, making it a potentially powerful imaging technique for studying thick, dose-sensitive specimens, showing a twofold dose advantage over EFTEM at 400 nm growing to fivefold at 800 nm.

Figure 4b compares the CTF for tcBF and DPC, which are the recorded signals we need for estimating the signal/noise ratios for the two methods. The detective quantum efficiencies (DQE) for tcBF and iDPC are proportional to squares of the CTFs plotted in Figure 4b (see online methods equation A16 for it is the DPC CTF and not the iDPC CTF that determines the iDPC DQE). While both approaches have the same information limit, tcBF has a higher information transfer at low spatial frequencies where much of the relevant structural information in a thick sample is located. In the language of ptychography^46^, tcBF is able to access both the double and triple-overlap regions, while DPC and single-sideband ptychography access only the double overlap (see online methods). tcBF is also able to surpass the real-space scanning Nyquist limit, offering a possibility for rapid data acquisition by trading detector pixels for real-space positions, important for out-running environmental noise in cryogenic experiments. Compared to ptychography, we find tcBF will still produce a robust image under thickness and dose conditions where our current ptychographic forward models fail to converge, and indeed there is a benefit to starting the ptychographic reconstruction from the information provided by tcBF, especially the estimate of the probe shape. Furthermore, at low doses where the signal is dominated by the central disk, our analysis summarized in Figure 4b,c gives insight into where the signals accessible to ptychography are encoded. Close to in-focus conditions, only the anti-Friedel term of the PCTF used in DPC and SSB imaging is available. At large defocus, the Friedel term of the PCTF used in tcBF and provides phase contrast at low frequencies is also accessible. This suggests low-dose pychography should be performed at large defocii conditions similar to those used for tcBF.

Overall, there are many lessons learned in the decades it took for EFTEM SPA to reach its present resolution that can also be applied to both the algorithm development and experimental design for tcBF and ptychography to boost their performances to comparable levels for thin specimens, and potentially well beyond for thick samples. One of the directions to improve is specimen motion correction. The movie mode and motion correction developed for conventional TEM operation mode effectively accounts for the thermal and mechanical drift and the beam-induced specimen motion^47,48^. In tcBF-STEM, upsampling effectively reduces the data acquisition time, thereby mitigating the impact of slow drift but beam-induced motion is not handled. Beam-induced motion, reflected by the large B factors in our tcBF reconstruction and other contemporary ptychography reconstructions, are probably the major factor limiting resolution. Current 4D-STEM pixel array detectors are still too slow to incorporate these corrections directly, but analogous correction modes should be possible. Both tcBF and ptychography already contain information in the overlapping probe positions that could be used to correct the beam-induced motion. At present this correction is limited by the dose/recorded diffraction pattern, but a fast detector design with a larger pixel count could address this by allowing for a larger illuminated area/pattern. In summaey, to fully exploit this information may require a new, faster generation of detectors and scan systems to meaningfully decipher the underlying specimen motion and time-ordered information. Future efforts aimed to enhance the performance of tcBF-STEM involve addressing beam-induced specimen motion and exploring the practical resolution limits of this technique. This includes using an increased probe convergence angle and higher-order probe aberrations, as well as exploiting the parallax effect to determine and correct the defocus for individual structures within thick sections.

### Online methods

#### tcBF-STEM upsampling

For the dataset shown in Fig.2, there are 256×256 scan positions with a 5.5 mrad convergence semi-angle probe-forming aperture (α). A scan step size of 8 Å is used, which sets a real-space Nyquist limit corresponding to 16 Å. With a defocus of 1.3 µm (nominal) is applied, the diameter of the illumination spot size on the sample plane is about 13 nm. With an 8-Å scan step size, the collected diffraction patterns contain a substantial amount of overlapping information. This surplus of information is utilized to achieve real-space upsampling through sub-(scan)-pixel image shifting. To implement upsampling, each image formed by a single detector pixel is padded before shift-correcting (Fig. S2), and then combined. The combined image is then weighted by the distribution of sub-pixel image shifts (Fig. S3). Different padding options are also compared and assessed in the supplemental information (Fig. S4). Zero-padding is observed to preserve information-transfer beyond the scan sampling. The PCTF simulation for tcBF and BF uses 5.5 mrad for convergence semi-angle and 700 nm defocus. Measuring image shifts in tcBF-STEM can also be regularized using the probe aberration function. All cryogenic tcBF images presented here benefit from this regularization, and a comparison with and without regularization is shown in (Fig. S9).

The limits of upsampling practically depends on several factors in addition to the optical resolution limit. Reciprocal-space sampling, real-space probe overlapping, and real-space image shift accuracy are critical factors for information retrieval through upsampling. Reciprocal-space sampling is primarily determined by the camera length, which is chosen to optimize the collection angle and angular resolution for a given detector. The degree of upsampling we can achieve is also limited by the accuracy of the image shift determination. Insufficient SNR in cross correlation can hinder the accuracy of image shift determination, which usually happens when the image SNR is low. It is possible to improve the accuracy by leveraging knowledge of the expected probe aberration function. It is also important to note that there is a trade-off between fineness of the reciprocal-space sampling and the SNR in the images formed by individual pixels in real space. Additionally, variations in the CTFs from higher-order aberrations and the impact of the aperture edge at different angular positions in the diffraction space, can also lead to false shift determinations.

#### The contrast transfer function for tilted-beam imaging

In linear imaging theory the image contrast *C* (*ω*)can be written as

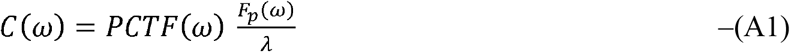

where *F*_*p*_ is the elastic scattering amplitude of the projected object and *λ* is the electron wavelength^48^. In general, the phase contrast transfer function (PCTF) can be complex, with the real part corresponding to angularly symmetric (i.e. Friedel-like) scattering, and the imaginary part to antisymmetric scattering. (At the lowest order of approximation, these terms would correspond to weak phase and weak amplitude approximations). Rose considered the phase contrast for samples which have undergone both elastic and inelastic scattering, with the case for a tilted beam given by equation 26 of reference ^48^. For weakly scattering objects, the quadratic and higher-order terms in his equation (26) can be neglected and a simpler, linear PCTF is given by Rose’s equation (33)

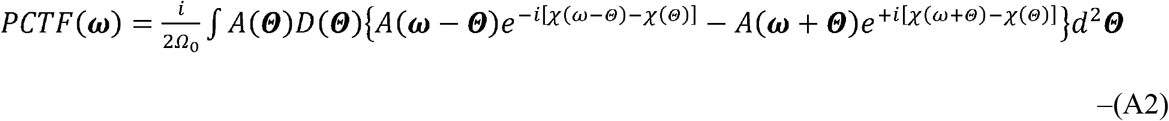

which we interpret here in terms of the STEM geometry where ***ω, Θ*** are momentum vectors projected onto the detector in the diffraction plane and normalized as scattering angles (which are a vector in this plane, hence the bold notation We also introduce the factor of ½ to be, consistent with the modern definition that the magnitude of the PCTF ≤ 1 is (See reference 48,2nd column top of pg259).*A* (*Θ*) and *D* (*Θ*) are the probe-forming and detector functions, which are 1 inside the apertures, and 0 outside. *χ*(*ω*)is the aberration function of the objective lens and Ω_0_≈ π α^2^ is the solid angle subtended by the objective aperture, which cuts off at angle α. For a pixelated detector with small pixels (i.e. the change χ in across a single pixel is small),*D* **(Θ)** ≈ δ (*Θ*) and equation (A2) simplifies to

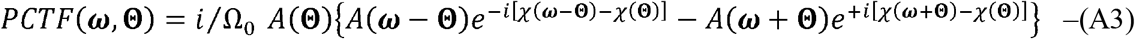

where ***ω***is the spatial frequency in the image, and ***Θ***is the collection angle (i.e. pixel) on the detector, *PCTF* (*ω* Θ)so gives the PCTF for the image formed by scanning the probe in sample plane and collecting the signal at pixel **Θ**on the detector. The *PCTF* (*ω* Θ)without an aperture is shown in Fig. S5 for a range of different tilts **Θ**and the corresponding (*ω* Θ) for an aperture is shown in Fig. S6.

For the special case of axial illumination (*Θ*=0) the PCTF reduces to the bright field CTF of−sin(χ(*ω*)). This would also have a cutoff at | ***ω*** |= α. The tilted beam case has non-zero contributions outside the aperture, up to a cutoff of 2αwhen | ***ω*** | = α from the terms *A* (*Θ*) *A* (*ω −Θ*) *A* (*Θ*) *A* (*ω −Θ*)This is the same information limit as the ADF and iDPC imaging, and double that of the axial bright field signal. The power spectrum of the apertured PCTF (Fig. S7) shows the double-resolution limit.

We can get a sense of how the aberrations lead to shifts in the image, by considering the special case where the dominant aberration is defocus so

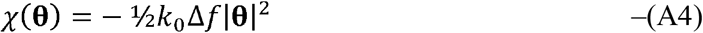

where 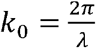. The PCTF then further simplifies to

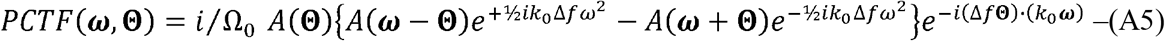

From the Fourier shift theorem, when transforming from the diffraction plane *k*_0_***ω***to the image plane ***x***, the 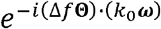 term in (A5) gives a shift of the image in real space of Δ*f* ***Θ***, i.e the shift proportional to the defocus and the angle from the axis on the detector. This is the tilt that is corrected by tcBF. The defocus aberration from the 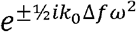 terms are still present in CTF and the tilt-corrected PCTF becomes

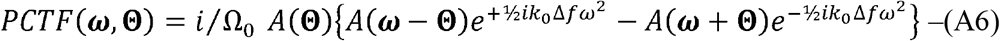

The tcBF CTF is obtained by summing over all tilt angles **Θ**. This is most easily accomplished by first summing symmetrically over pairs of angles at **Θ**and − **Θ**:

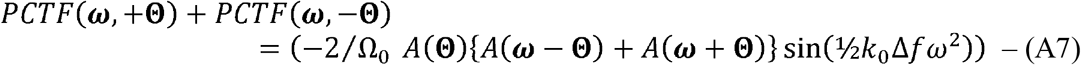

and then completing the sum over half of the central disk (say all **Θ***x* >0). In polar coordinates **Θ**=**(Θ**, ϕ)and for a disk of diameter α we integrate over **Θ** and ϕ

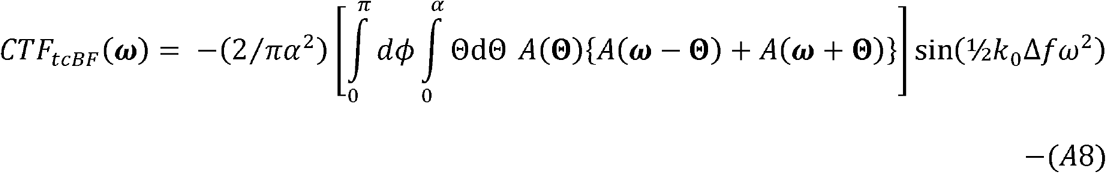

The integral over **Θ**gives the area of the overlap of disks of diameter α that are ***ω*** apart, and can be found in the appendix of reference 48 as

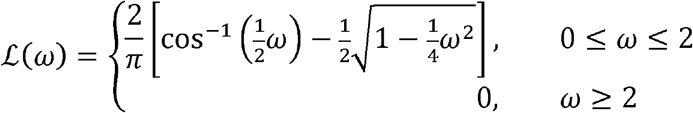

ℒ (*ω*)is the well-known envelope for a self-luminous object, such as for the annular dark field contrast transfer function. The tcBF CTF can then be written more compactly as

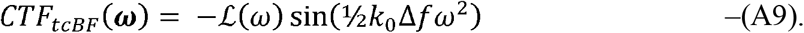

### Comparison of the contrast transfer function for tcBF with DPC

The optimal CTF for DPC and iDPC is the in-focus condition with no aberrations inside the aperture. Then *χ*(**θ**)= 0 and the general PCTF simplifies to

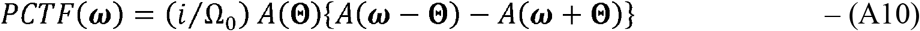

i.e ℜe (*PCTF*) = 0 at zero defocus, and only the antisymmetric component remains (Fig S6c). The DPC_x_ signal is produced by subtracting all the left-tilted (**Θ**_*x*_<0) from the right-tilted **(Θ**_*x*_<0) detector signals and then summing to produce the DPC CTF of Figure 4b.

At **Θ** = 0 the, *PCTF*(***ω***)=0 and the PCTF remains 0 so long as |***ω*** |< *α*, | ***ω*** − **Θ** |< αand | ***ω***+ **Θ** |< *α*.giving the white regions in each frame of Fig. S6c. In ptychography, this is referred to as the triple overlap region^46^, reflecting the simultaneous overlap of the +*ω* and −*ω* beams with the incident beam (in ptychography, this is usually displayed in detector plane **Θ**for a range of selected ***ω*** while we have displayed the ***ω*** plane for a range of selected **Θ**), and is zero for in-focus imaging. When a phase shift is deliberately introduced, this triple-overlap provides the phase contrast for BF imaging, but still remains zero for DPC and single-side band (SSB) ptychography (white regions of Fig S6b). DPC and SSB rely on the double-overlap region where |***ω***| < *α* and either |***ω*** − **Θ** |<α or |***ω****+* **Θ**| *<α*, but not both. Again, the information limit is the largest value of w for which the PCTF is non-zero. This occurs at |**Θ** | = *α* and *ω*= 2, so the largest non-zero value of |***ω***| is *ω*= 2*α*, double the radius of the aperture.

It is important to note that the CTF in the triple-overlap region has double the amplitude of that of the double-overlap region(Figure 3 of reference ^46^). This suggests that tcBF should have the potential to reach double the dose-efficiency of DPC at below spatial frequencies where |***ω***| < *α* and a *π*/2 phase shift can be introduced through the aberration function. This difference becomes very noticeable at low spatial frequencies where the double overlap terms tend to zero, and the triple overlap contrast can be boosted by increasing defocus.

**Figure 3.**
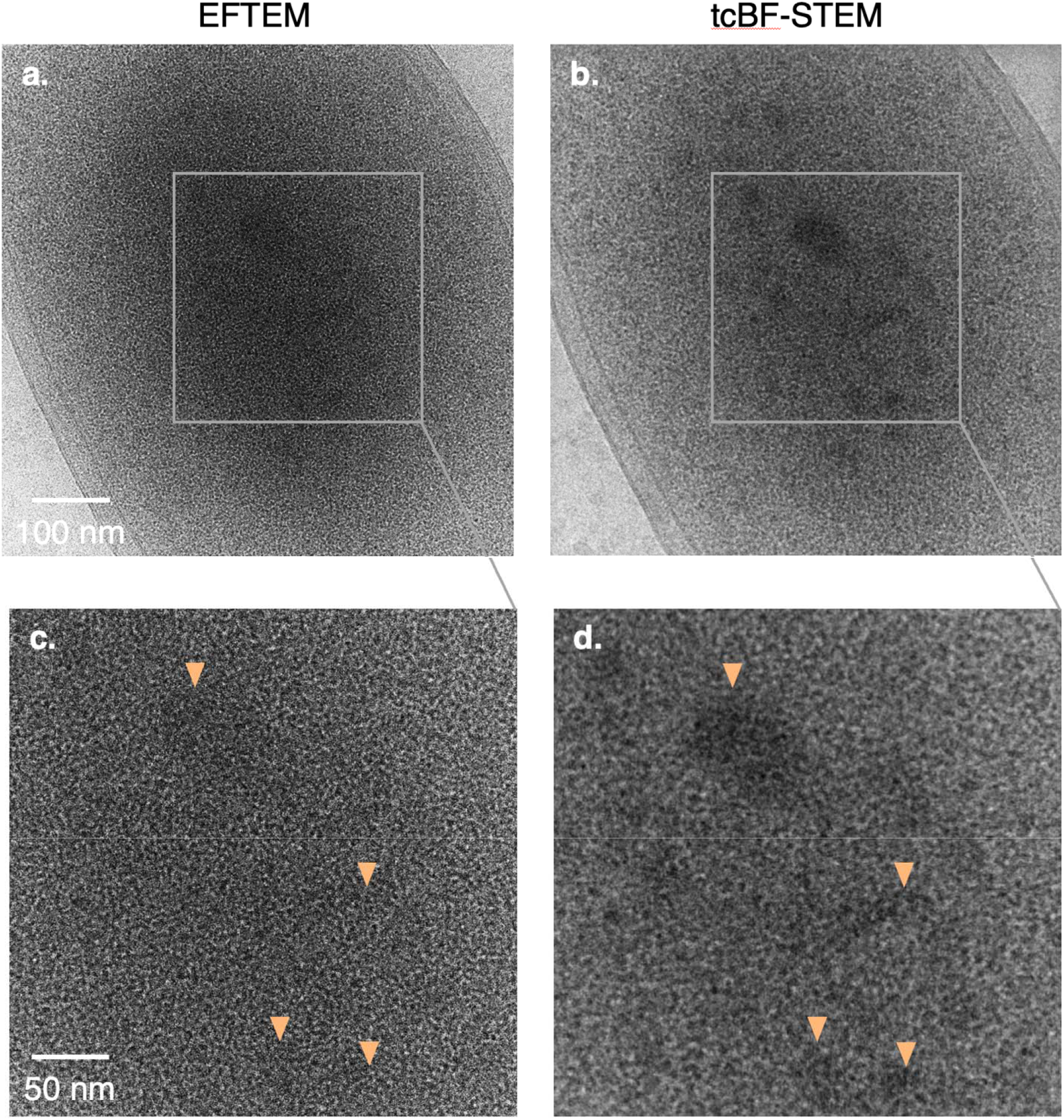
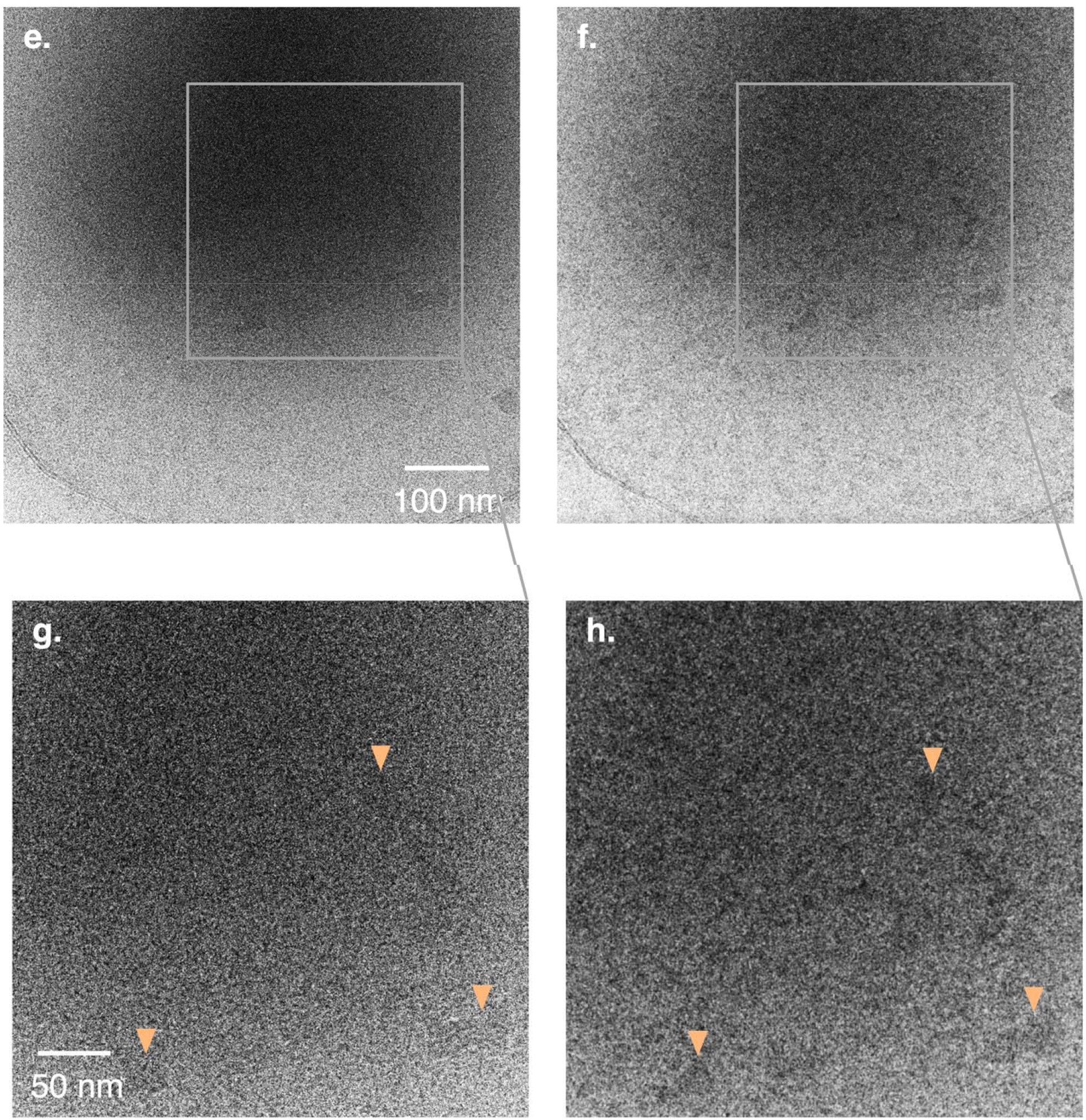
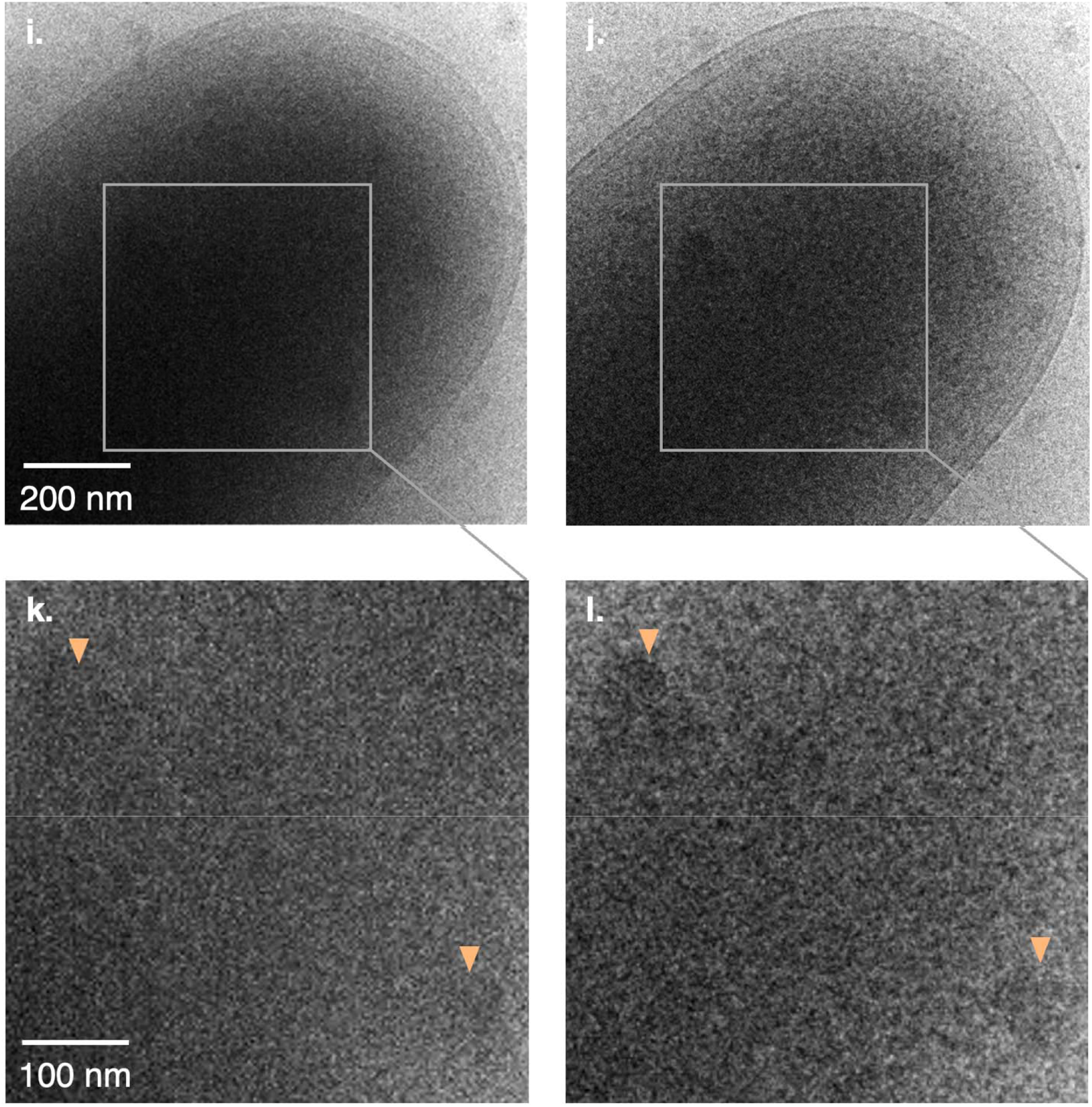
Comparison of EFTEM and tcBF-STEM with different doses, defoci and acquisition orders. The same region of interest in various specimens was imaged successively in order to compare the techniques. For each comparison, the total dose measured over vacuum and the electron acceleration (300 kV) are the same, and the average thickness can be estimated with the EFTEM images using the ratio of I0/I and the inelastic MFP, similar to Fig. S1. The dose efficiency of the two techniques is compared by the ratio of remaining electrons in the images to the incident total electrons. Overall, for the samples demonstrated here, tcBF is observed 3-3.5x higher collection efficiency than EFTEM at a similar incident dose/unit area. For EFTEM images, slit widths are all 10 eV and defoci are measured with CTFFIND4^42^. For tcBF images, defoci are measured with the image shifts. (a) and (b) are EFTEM and tcBF images of an intact *E*.*coli* cell. With tcBF (d), features in the interior region of the cell are effectively resolved, whereas in EFTEM (c), although the same features are discernible, they are less visible. In (e) and (f), images with EFTEM and tcBF of a vesicle at similar measured defoci are shown. Again, in the thick region tcBF reveals clearer features compared to EFTEM. For another comparison on *E*.*coli* at a low dose of 0.5 e^-^/A□^2^ (i-l), tcBF is able to resolve features that are otherwise indiscernible with EFTEM. Thickness and other experimental details in Table I.

### Comparison of the Detective Quantum Efficiency (DQE) for tcBF and iDPC

The iDPC CTF is obtained from the DPC CTF by integration in real space, corresponding to a division by spatial frequency in Fourier space as

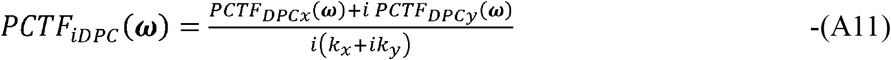

The power spectrum of the recorded DPC image in the presence of a noise spectrum N(ω) is

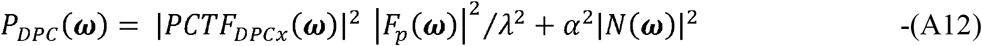

The power spectrum for iDPC based on the DPC measurement with noise is

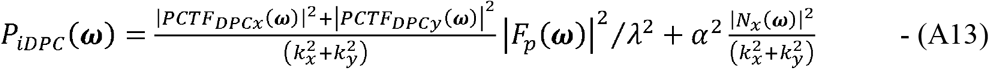

The DQE of the measurement with a noise power spectrum, *NPS*(*ω*), is

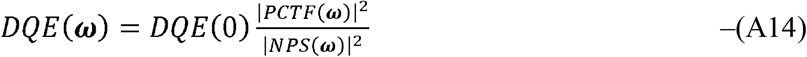

For an ideal detector pixel, *DQE(0)* =1 and for DPC imaging the *DQE(ω)* becomes

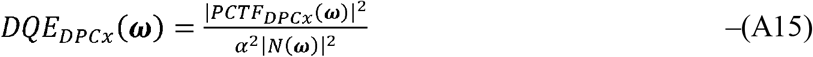

and after integrating, the DQE for iDPC becomes

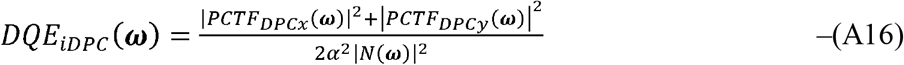

This has a very similar shape to the DPC DQE since the noise is amplified in the same way as the signal.

Similarly, applying equation A10 for tcBF we find the tcBF DQE to be,

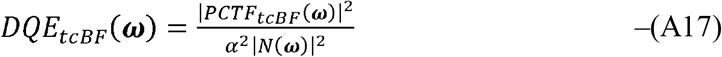

For an ideal detector the noise spectrum is only from Poisson noise, which is flat, so the differences in DQE for tcBF and iDPC can understood by comparing the squares of the PCTFs for tcBF and DPC (not iDPC). These are shown in figure 4b. As a consequence, iDPC has a poor DQE at low spatial frequencies compared to tcBF.

**Figure 4.**
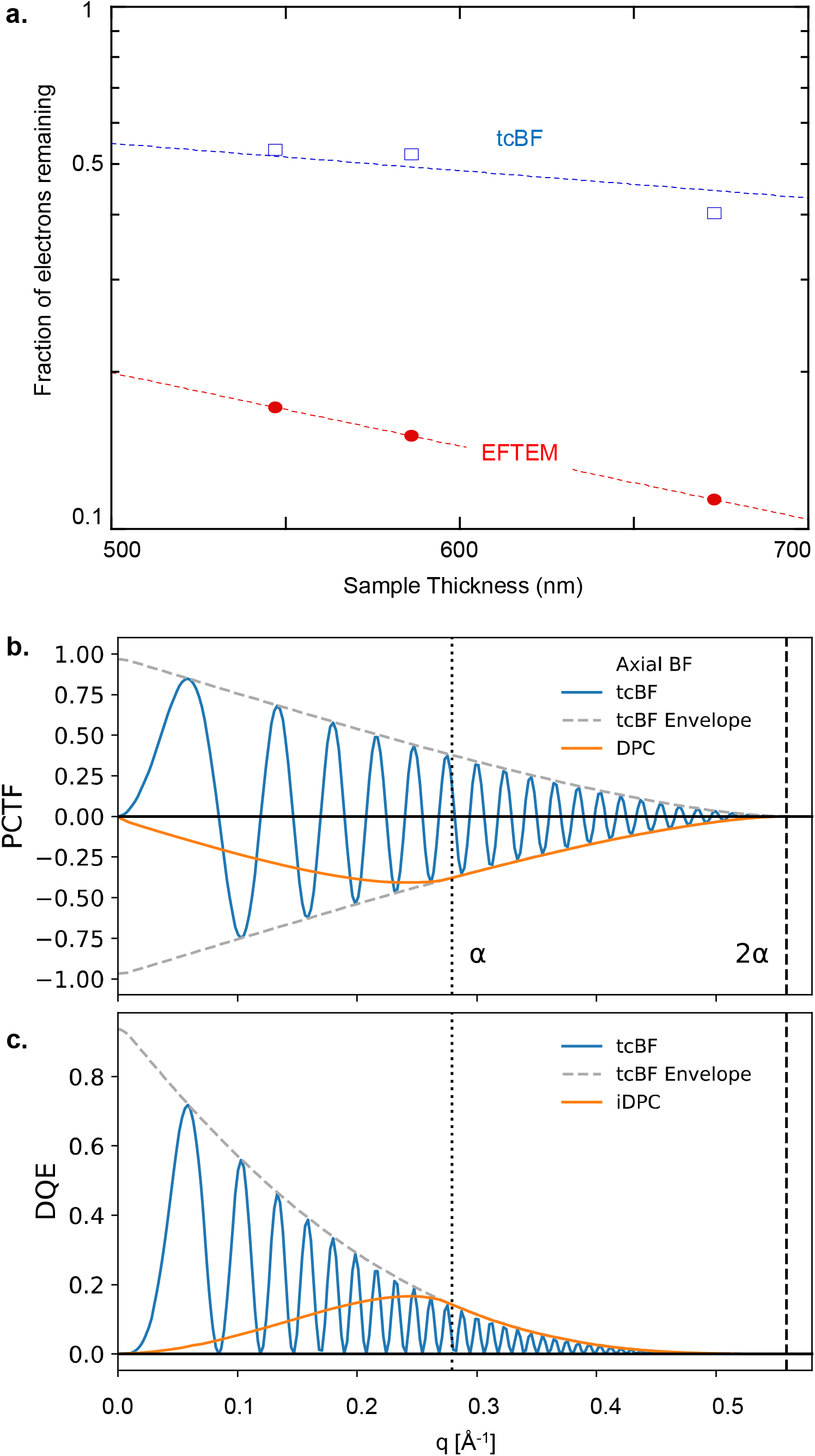
(a) Collection efficiency of EFTEM (no objective aperture) and tcBF (7 mrad aperture), showing the measured fraction of electrons left in the image compared to the incident beam as a function of sample thickness from the data sets in table 1. tcBF is seen to retain over 3-4x more signal than EFTEM. The sample thickness is determined from the EFTEM fraction, assuming an inelastic MFP of 310 nm. From this, the decay of the unfiltered tcBF images gives an elastic MFP of 830±50 nm. (b) Comparison of the contrast transfer functions for tcBF, axial BF, and in-focus DPC for a 5.5 mrad probe-forming aperture, a. The axial BF CTF cuts off at a, while the DPC and tcBF information limits extend to 2a. The damping envelope for tcBF follows the classic double-overlap form expected from summing over the tilted CTF functions in supplementary figures S6 and S7. The DPC signal peaks close to a and is suppressed at low frequencies compared to the defocus-optimized tcBF but is more efficient from a to 2a. The iDPC CTF has the same shape as the damping envelope but does not reflect the true information transfer. The iDPC CTF is obtained by dividing the measured DPC signal by spatial frequency, which also amplifies noise by the same proportion, resulting in a vanishingly small signal/noise ratio at low spatial frequencies (see online methods for analytic derivations). (c) The result is the DQE for iDPC is the same as that for DPC and both have poor efficiency at transferring low frequencies. Defocused tcBF is very efficient at transferring low frequencies.

**Figure 5.**
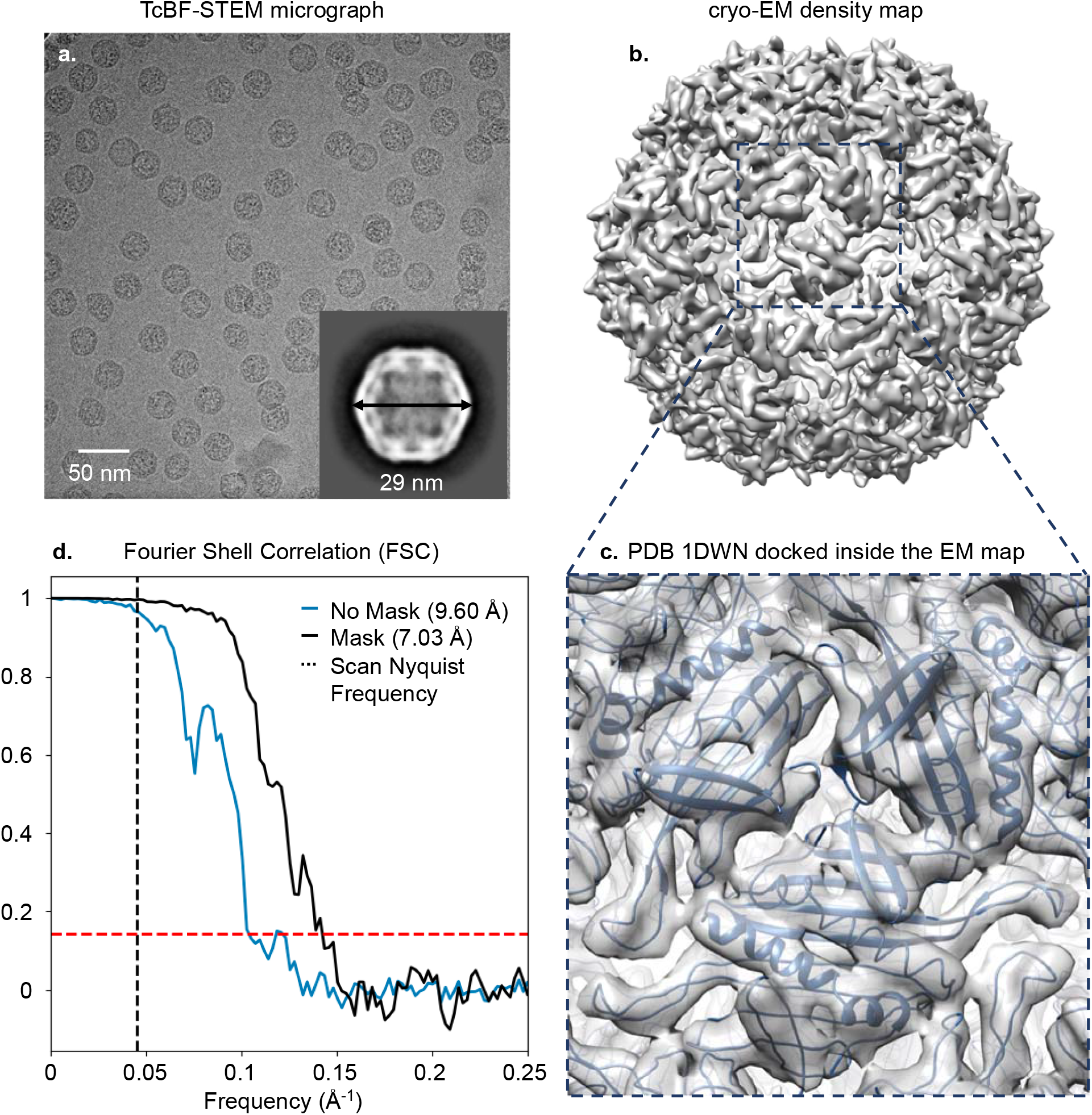
Single particle analysis 3D reconstruction from tcBF-STEM imaging of hydrated vitrified coat protein of bacteriophage PP7. A representative up-sampled tcBF-STEM image at 300 kV with 11 A□ scan step size and a total dose of 45 e-/A□^2^ is shown in (a) with a 2D class average in the inset. The 3D density map resolved from 789 particles is shown in (b) with a zoom-in view of the PDB 1DWN model fit inside the density map in (c). ∼7 A□ nominal resolution is reached based on the Fourier Shell Correlation (FSC) with 0.143 cutoff.

### Mean Free Paths and Thickness Estimates

Measurements of the inelastic MFP (scaled to 300 keV^49^) range from 100 nm for amorphous carbon to 275 nm for proteins to 310 nm for vitreous ice^50^, scaling roughly with the degree of hydrogenation. For our thickness measurements we use the inelastic MFP of ice. The elastic MFP is more strongly dependent on the range of collection angles as the elastic scattering has a much wider angular distribution than inelastic scattering. Thus, what is often reported is n, the ratio of elastic to inelastic scattering for a given measurement geometry, and this is in the range 2-5, with 3 being a typical value for cryoEM of organic systems^51^, suggesting a typical elastic MFP is about 700-900 nm. We calculated the elastic MFP from a multislice simulation of amorphous ice, for a 50 and 200 nm thick supercell and a 5.5 mrad convergence and collection angle at 300 keV. Averaging over multiple configurations, we fit the decay of the central beam to find the elastic MFP, *λ*_el_ = 774± 45 nm. The elastic MFP sets a thickness for which the dominant contrast mechanism crosses over from phase contrast to scattering absorption contrast. In relating the signal remaining in an energy filtered image, *I*_*EFTEM*_(*t*)*=I*_*TEM*_ *exp*(− *t / λ*_*in*_) and *I*_*TEM*_ is the corresponding unfiltered image. Even when no objective aperture is used, there is still some high-angle elastic scattering (including backscattering) that does not reach the detector, so not all of the incident beam is collected and *I*_*TEM*_(*t*)=*I*_0_exp(−t/*λ*,_*HA*_). Combining these results, we get

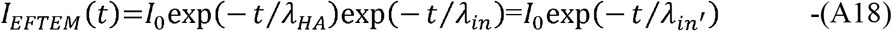

For our microscope, we measured, *λ*_*HA*_ = (46 ± 1), *λ*_*in*_, and it is convenient to keep the functional form, *λ*_*HA*_ = α*λ*_*in*_ From eqn (A14) we can calculate the high angle correction to the inelastic mean free path as

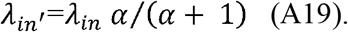

so, *λ*_*in*_^′^=0.979, *λ*_*in*_ = 303 nm for ice.

### Comparative analysis on thick samples

The organelles shown in Fig. 1i-n and Fi. 3e-h were isolated and purified from the HEK293T cells. Cells were mechanically lysed by osmotic shock and needle shearing^52^. The STEM images were recorded using an EMPAD^19^ on a TFS Krios G4 with a 7 mrad semi-convergence angle and a 2.8nm scan step size. 256*256 scan positions were collected. The corresponding EFTEM images were recorded using a Falcon 4i detector and the Selectris X energy filter with a slit width of 10 eV on the same TFS Krios G4. Acceleration voltage was 300 kV and the spherical aberration of the objective lens was 2.7 mm. tcBF images are reconstructed with the iterative alignment provided in py4DSTEM^32^ and upsampling is implemented in an in-house Python package based on the method described in the upsampling section.

The E. coli specimen shown in Fig. 3 were prepared from the GL002 strain and plunge frozen with 200-mesh Quantifoil 2/1 holey carbon copper TEM grids. For Fig. 3i-l, the images were recorded on a customized Thermo Scientific Titan Themis with Gatan 626 cryo-transfer holder at 300 kV. The STEM images were recorded using an EMPAD^20^ with 2 mrad convergence angle. tcBF images were reconstructed with in-house Python package where the alignment algorithm is based on rigid shift registration between every possible pair^53^ and upsampling algorithm as described in the section. The EFTEM images were acquired with a K2 Summit direct detector (Gatan) operating in linear mode. For all the EFTEM images, short exposures were collected in the movie mode and cross-correlated with the number of frames chosen to match the dose of the corresponding STEM image.

### Single particle analysis on VLPs

The specimen is a coat protein of bacteriophage PP7 self-assembled during recombinant expression in E. coli. TEM grids used are R1.2/1.3 mesh 300 UltrAuFoil. STEM images were acquired on the customized Thermo Scientific Titan Themis with a Gatan 626 cryo-transfer holder. Images were recorded with 300 kV acceleration voltage on an EMPAD-G2 detector^20^, 8 mrad semi-convergence angle, 11 Å scan step size, 45 e^-^/Å^2^ total exposure dose and upsampled to an image pixel size of 2.77 Å. A typical scan size is 512*512 with 100 µs dwell time. tcBF images are reconstructed with the iterative alignment provided in py4DSTEM^32^ and upsampling is implemented in an in-house Python package based on the method described in the upsampling section. The SPA reconstruction is obtained with cryoSPARC^44^ where CTFFIND4^42^ is used to estimate global CTF. Particles are picked with a template generated by manually-picked particles. The final 3D reconstruction has icosahedral symmetry and a dynamic mask imposed. The comparative analysis of the VLPs with EFTEM is performed with a Cs-corrected TFS Krios, shown in Fig. S11. The EFTEM data were acquired at an accelerating voltage of 300 kV, a pixel size of 1.076 Å and a total dose is 52.18 e^−^/Å^2^. The defocus ranges from -1µm to -2µm. The final reconstruction is obtained with 900 particles and the analysis is also done with cryoSPARC^42^.

## Acknowledgements

This work is supported by NSF (DMR-1654596, DMR-1429155, DMR-1719875, DMR-2039380), the Packard Foundation, and Chan Zuckerberg Institute for Advanced Biological Imaging. This work made use of the instruments at Chan Zuckerberg Institute for Advanced Biological Imaging, the Cornell Center for Materials Research (CCMR) Shared Facilities and PARADIM. CCMR facilities and X.S.Z. are supported through the NSF MRSEC program (DMR-1719875). PARADIM and S.E.Z. are supported by the NSF MIP program (DMR-2039380). We are grateful for all the time that Lena was able to share with us. May her memory be a blessing. The authors thank Dr. Tianhong for inspiring discussions on tcBF upsampling. The authors appreciate Dr. Yasu Xu for providing the E.coli specimens, and Dr. Manuel D. Leonetti’s group for providing the cell lines for the organelle specimens. In addition, the authors want to thank Dr. Earl J. Kirkland for helpful discussions on tilted BF CTFs and Paul Cueva (NSF PHY-1549132) for help with aberration and tilt measurements in 4D-STEM. The authors acknowledge Dr. Georgios Varnavides, Dr. Stephanie M. Ribet, and Dr. Colin Ophus for helpful discussions on improving algorithms for tcBF. The authors also thank Dr. Bridget Carragher, Dr. Clinton S. Potter, and Dr. David Agard for advice on experimental designs for comparing tcBF-STEM to EFTEM, as well as for insights on future steps to improve the technique.

## Author Contributions

Y.Y., K.A.S., D.A.M., and L.F.K. designed the tcBF experiments. Y.Y, K.A.S., and K.X.N. performed the tcBF experiments. Y.Y., K.A.S., M.C., K.X.N, D.A.M. and L.F.K. developed tcBF algorithm and analyzed the tcBF data. S.E.Z. and D.A.M. calculated the tcBF CTFs. R.P. D.S. and H.S. prepared the purified cellular organelle samples. Y.Y., R.P., M. K. and C.D. analyzed the single particle data. X.Z. and Y.Y. performed iDPC experiments. X.Z. analyzed iDPC data. Y.Y., K.A.S., L.F.K., and D.A.M. wrote the manuscript with input from all authors.

## Data Availability

The 4D-STEM data sets for Fig. 1l-n and Fig.2a-h are available on Zenodo (DOI: 10.5281/zenodo.10825339), along with the corresponding EFTEM images.

## Code Availability

House built python packages for tcBF-STEM are available on Github at https://github.com/yyu2017/tcBFSTEM.

## Figures

**Fig. S1.**
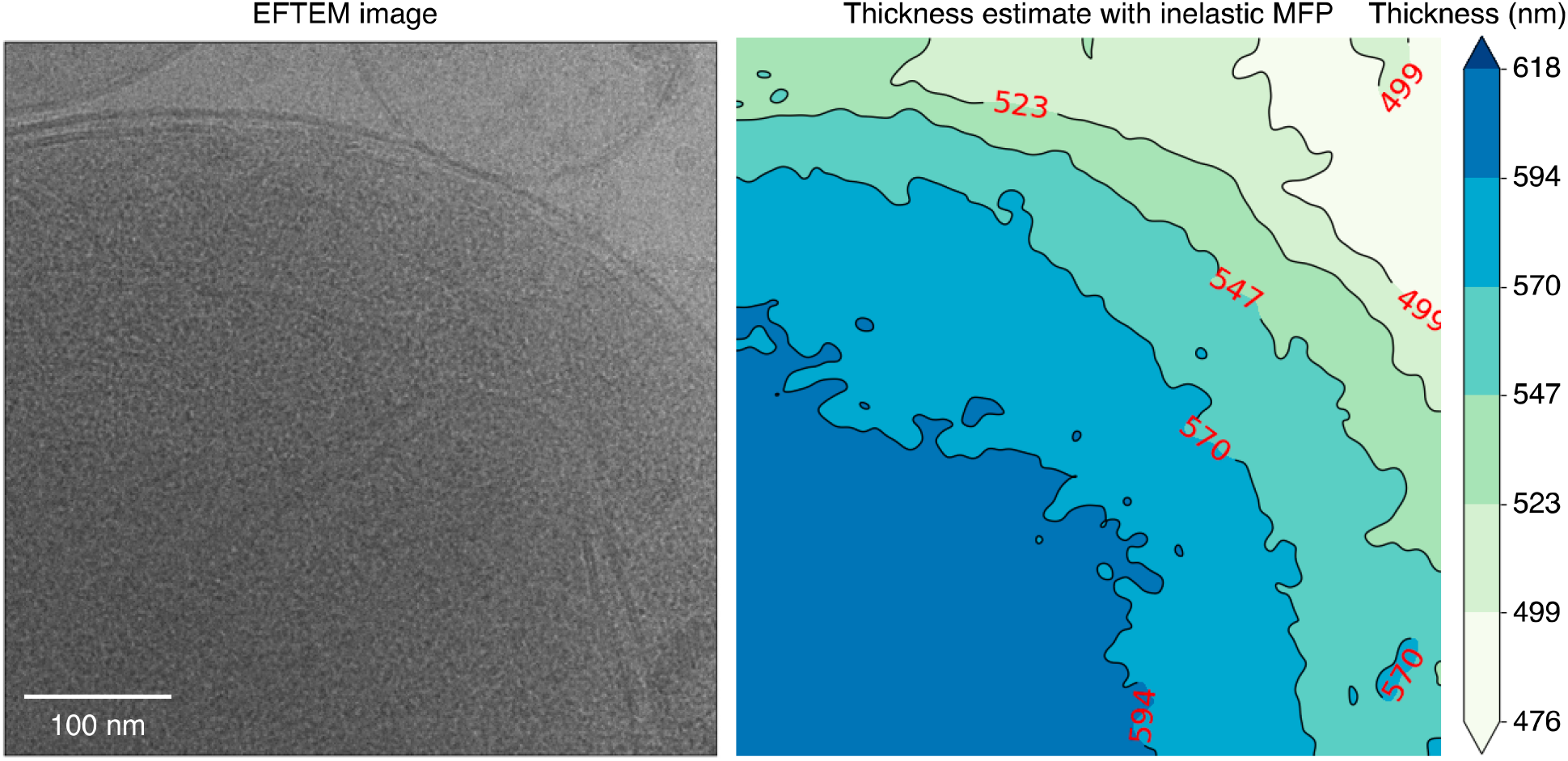
A contour plot of the thickness estimate for the sample shown in Fig.1i-n. The thickness is estimated with the inelastic MFP using Beer’s law. Using the EFTEM dataset, we obtain a ratio of I0/I, where I0 is the intensity recorded over vacuum, and I is the energy filtered intensity with a 10-eV slit recorded over the sample. The thickness is estimated ln(I0/I)*inelastic MFP. The inelastic MFP used here is 310 nm for vitrified ice at 300 kV^41^.

**Fig. S2.**
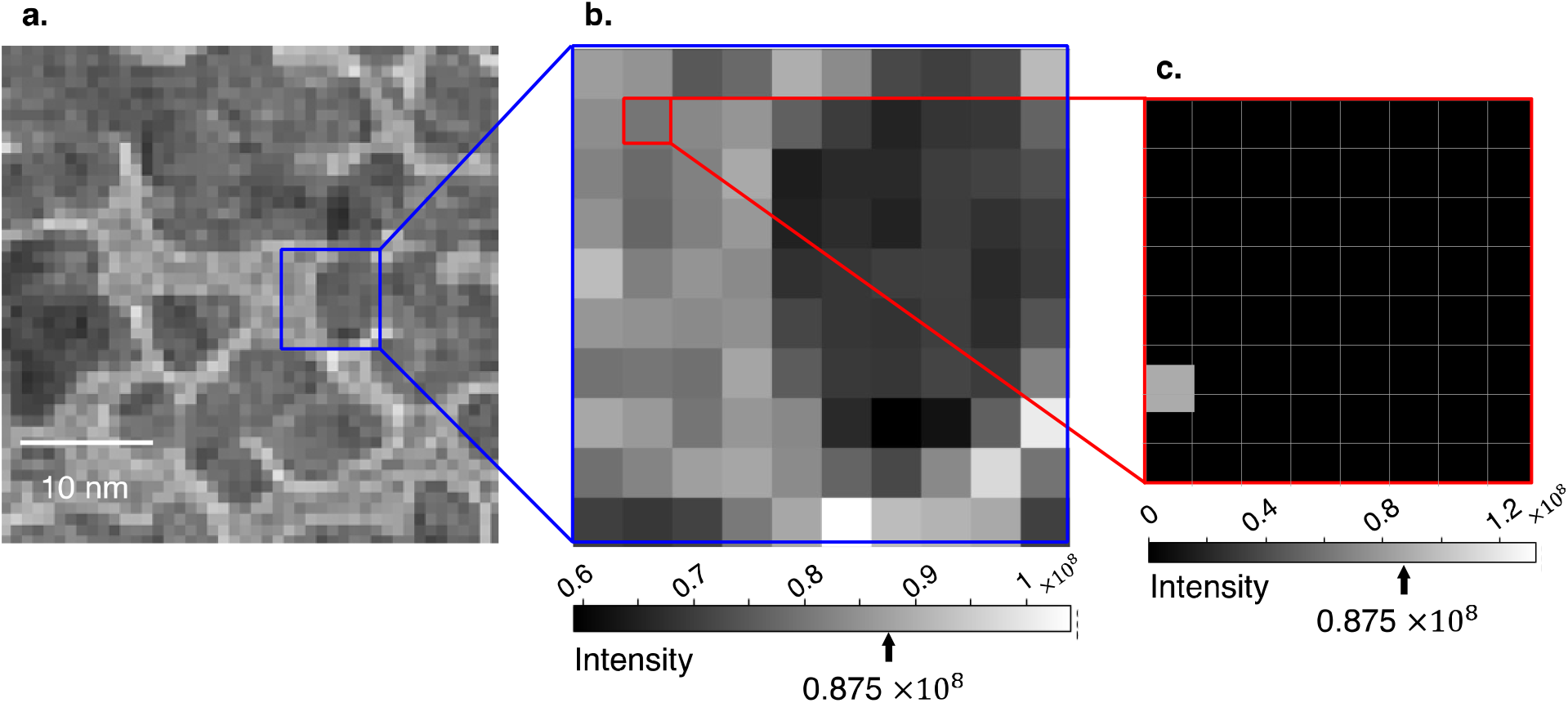
To facilitate sub-scan-pixel image shifting, each real-space image formed by a single detector pixel is first padded with zero-intensity pixels. In (a), there is a bright field image formed using a single detector pixel for a standard gold-on-carbon sample under the previously described imaging condition in Fig.2. Each pixel (b) in the image (a) becomes an 8×8 pixel block (c) after padding. The padding process only involves inserting zero values between the original pixels without altering their values.

**Fig. S3.**
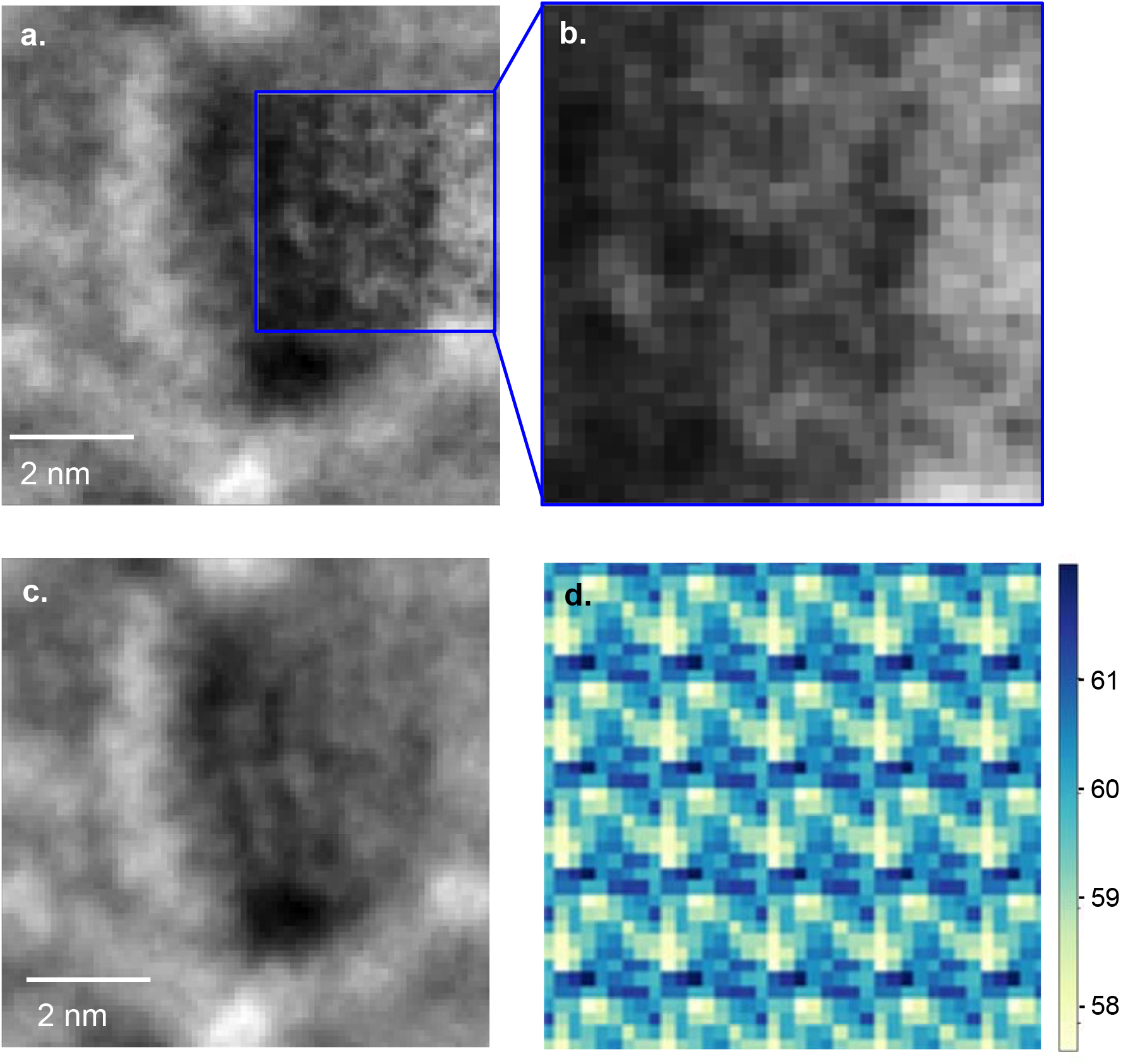
Normalization of uneven distributions of sub-pixel shifts: the same up-sampled image shown in Fig. 2d with the periodic intensity variations amplified in the blue-boxed area (b) for better visibility. By tracking the sub-pixel shift distribution (d) and applying an intensity normalization based on this distribution, the periodic artifacts can be corrected (c).

**Fig. S4.**
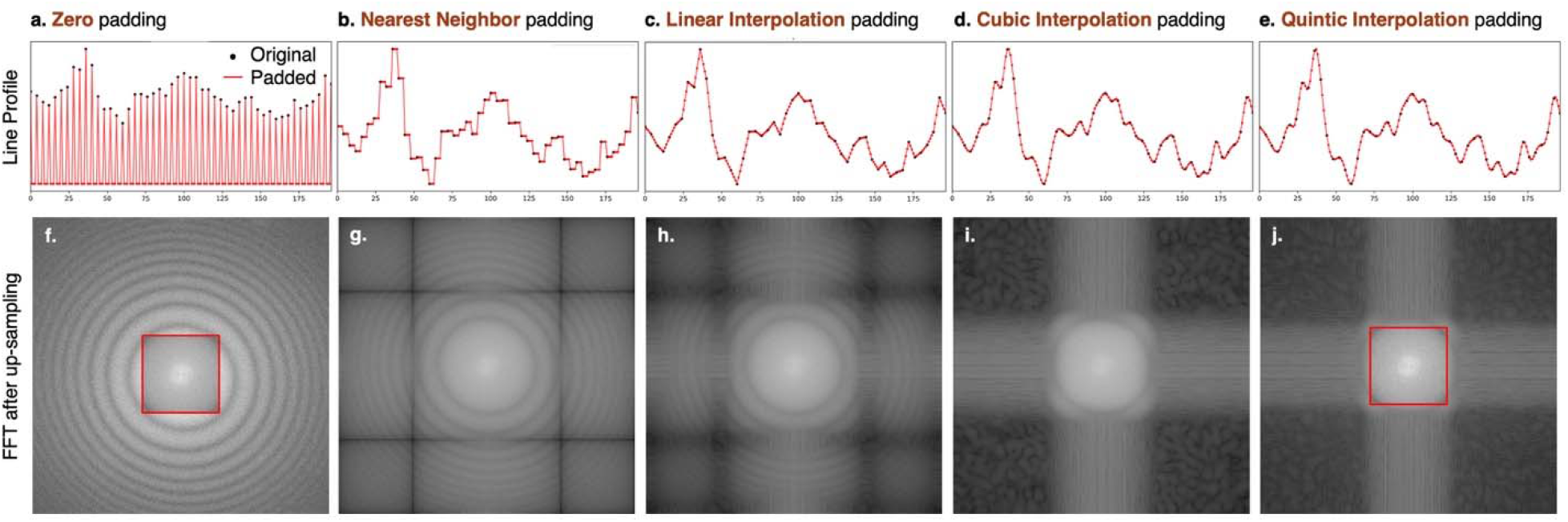
Higher-order interpolation padding: line profiles (left) illustrate in 1D the results of padding with zero-intensity pixels and progressing to quintic interpolation values. The corresponding FFTs of the up-sampled images are shown on the right.

**Fig. S5.**
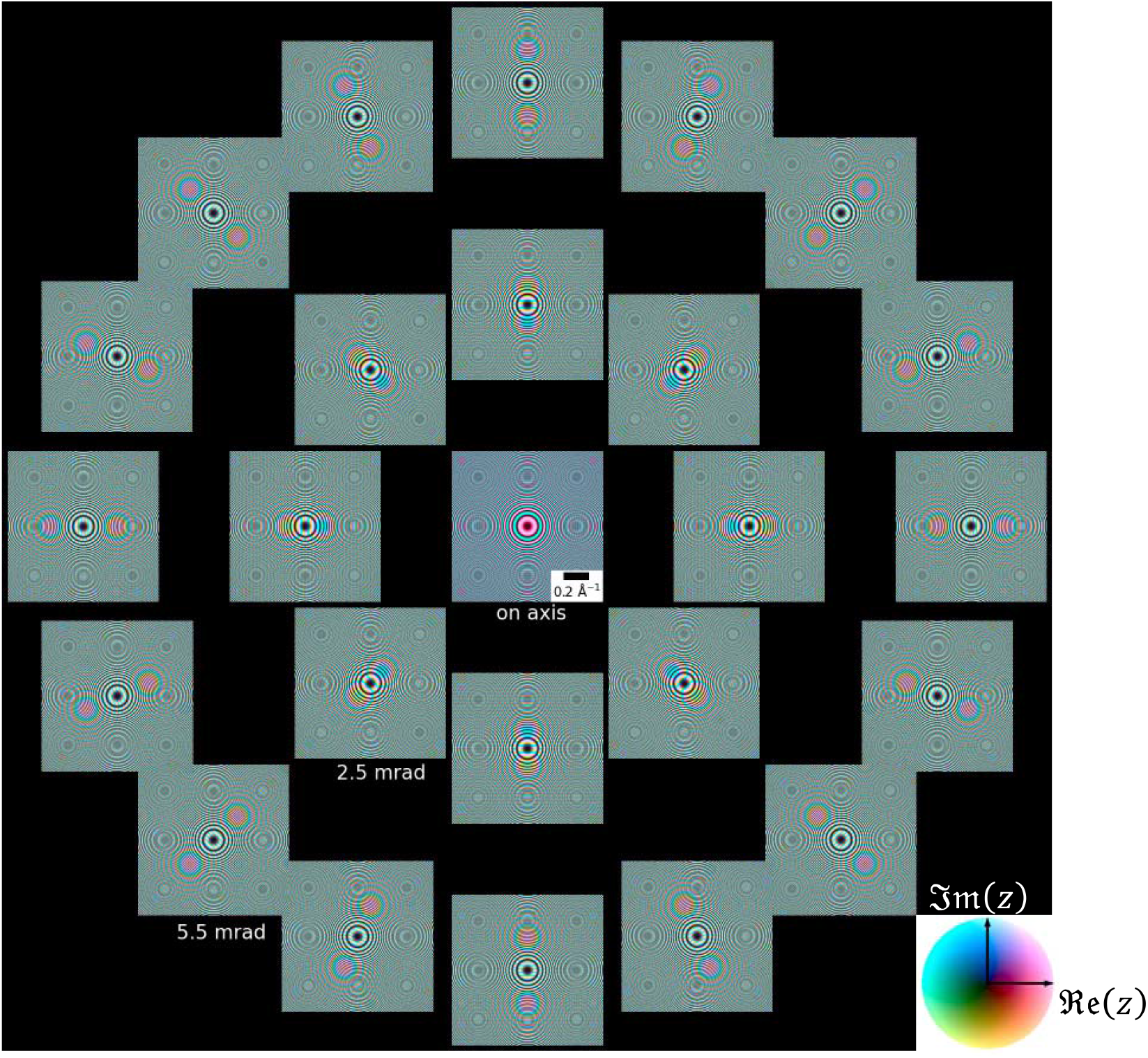
The aperture-free, complex PCTF for 300 kV electrons and 700 nm defocus in a Zemlin tilt-tableau out to 5.5 mrad of tilt. The x-y coordinates within each frame are spatial frequencies of the image, w, and the tilt offset of each frame is 0.With no objective (condenser) aperture and no higher order aberrations, the power spectrum of each image under tilted illumination is identical.

**Fig. S6.**
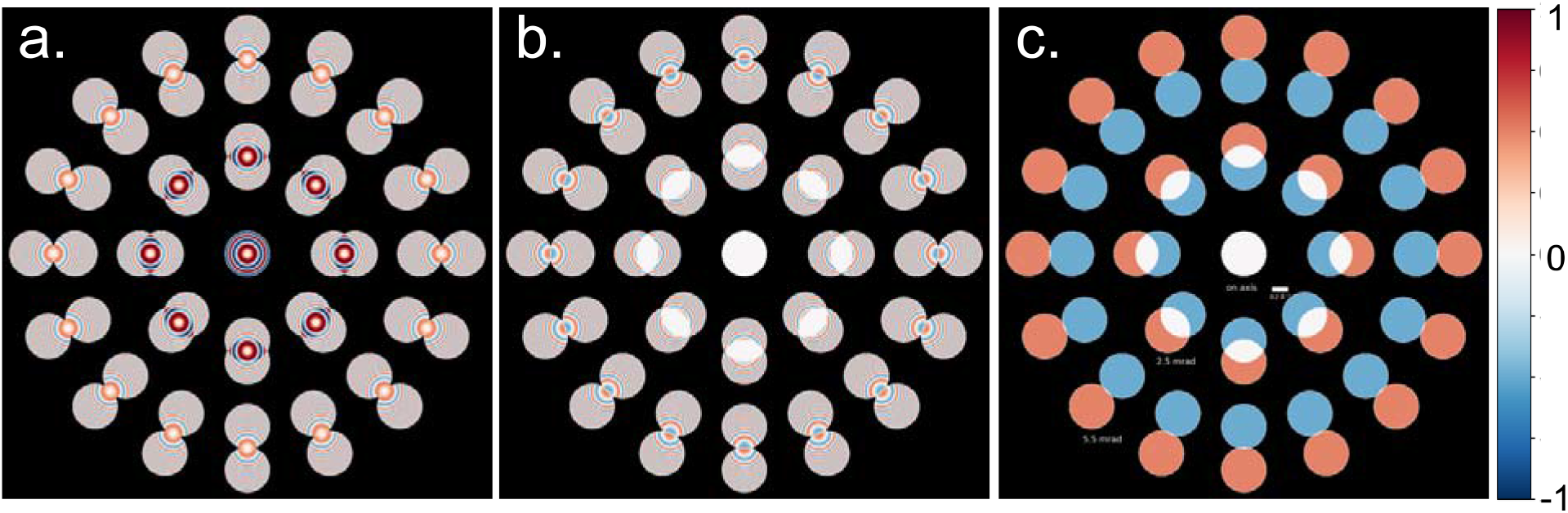
The symmetric and antisymmetric components of the PCTF for 300 kV electrons with a 5.5 mrad objective (condenser for STEM) in a Zemlin tilt-tableau out to 5.5 mrad of tilt. (*a*) ℜ 𝔢 (*PCTF*) at 700 nm defocus showing the symmetric, Friedel term, (b) 𝔉 𝔪 (PCTF) at 700 nm defocus showing the anti-symmetric, anti-Friedel term. The phase ramp across each individual PCTF reflects a shift in real space of the imaged object. Shifting the individual images corrects for the tilt-induced phase ramp, and subsequently summing the tilt-corrected images gives the PCTF shown in Figure 2f. (c) 𝔉 𝔪 (*PCTF*) at zero defocus, again showing its anti-symmetric nature. The DPC-x image is formed by subtracting the left tilts from the right.

**Fig. S7.**
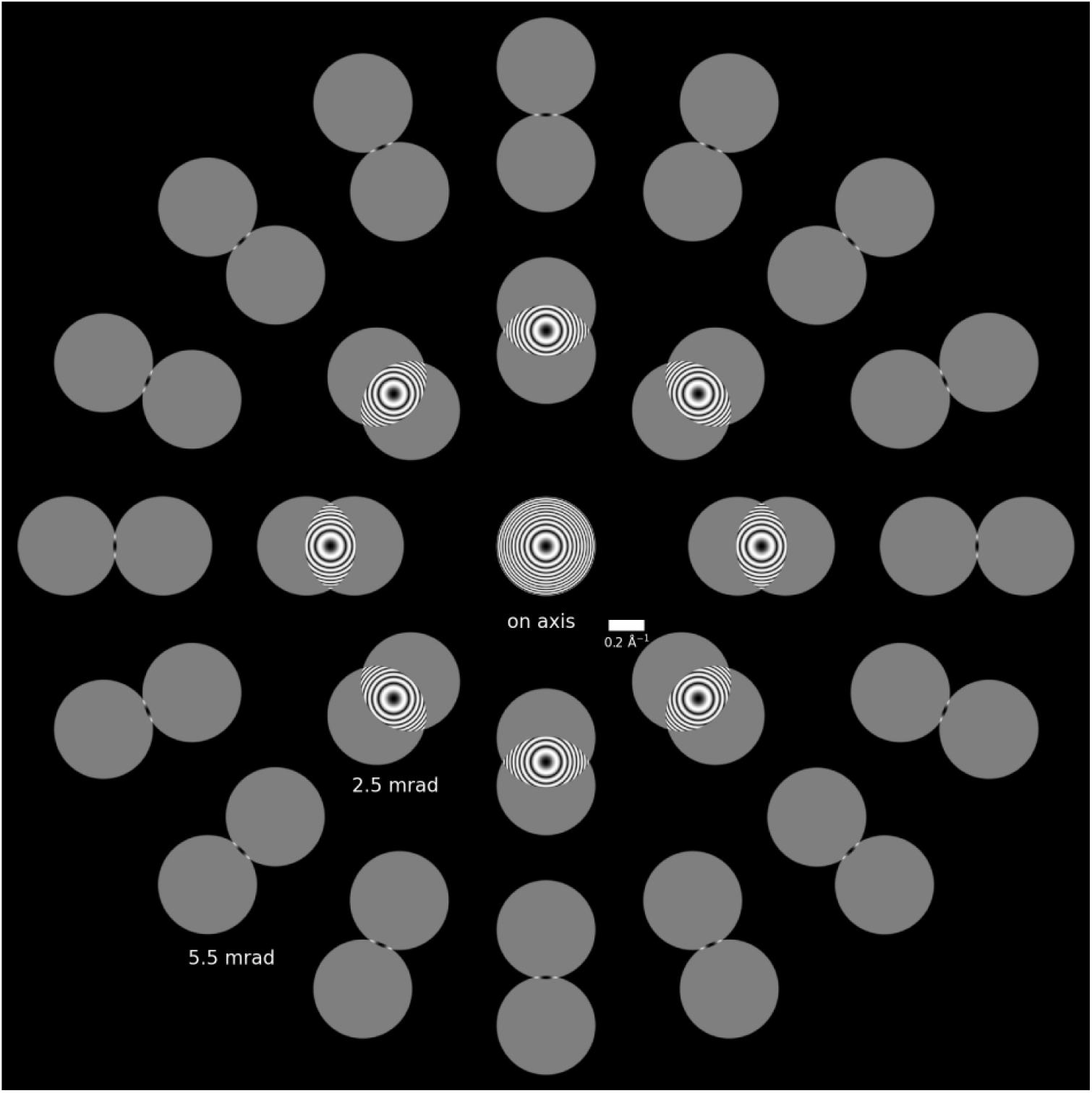
The power spectrum for 300 kV electrons and 700 nm defocus with a 5.5 mrad objective (condenser) in a Zemlin tilt-tableau out to 5.5 mrad of tilt. This highlights the strong modulations in the overlap region, and the weaker transfer in the sidelobes, but with an information limit of twice the aperture radius.

**Fig. S8.**
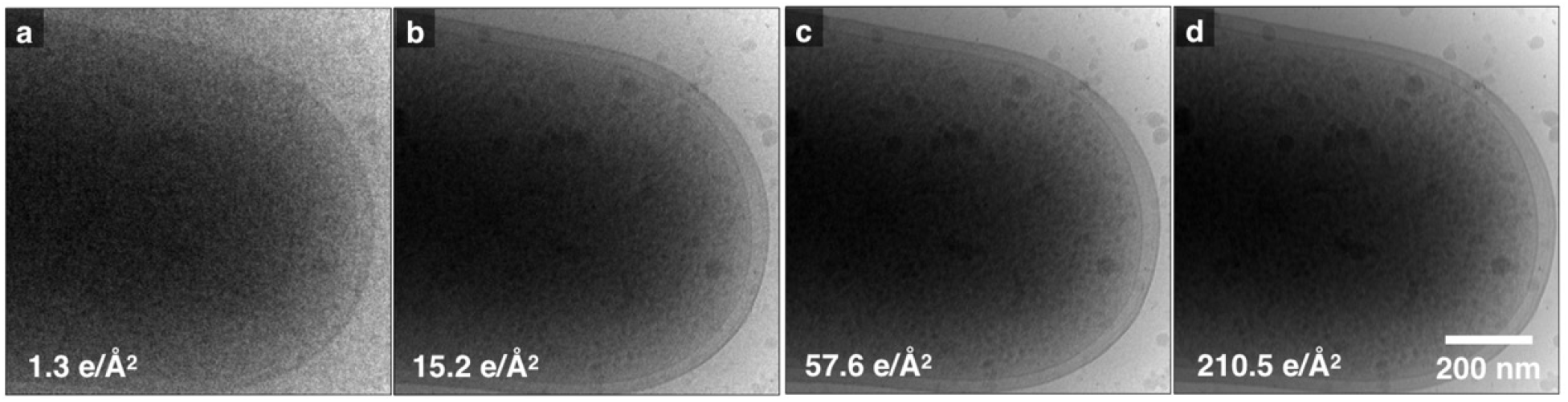
Dose Tolerance for tcBF-STEM images collected sequentially with increasing dose: (a) 1.3 e^-^/Å^2^, (b) 15.2 e^-^/Å^2^, (c) 57.6 e^-^/Å^2^, (d) 210.5 e^-^/Å^2^. Large-length-scale features in the specimen appear tolerant to a high cumulative dose, with no bubbling appearing even in the final exposure where the cumulative dose is 286 e^-^/Å^2^.

**Fig. S9.**
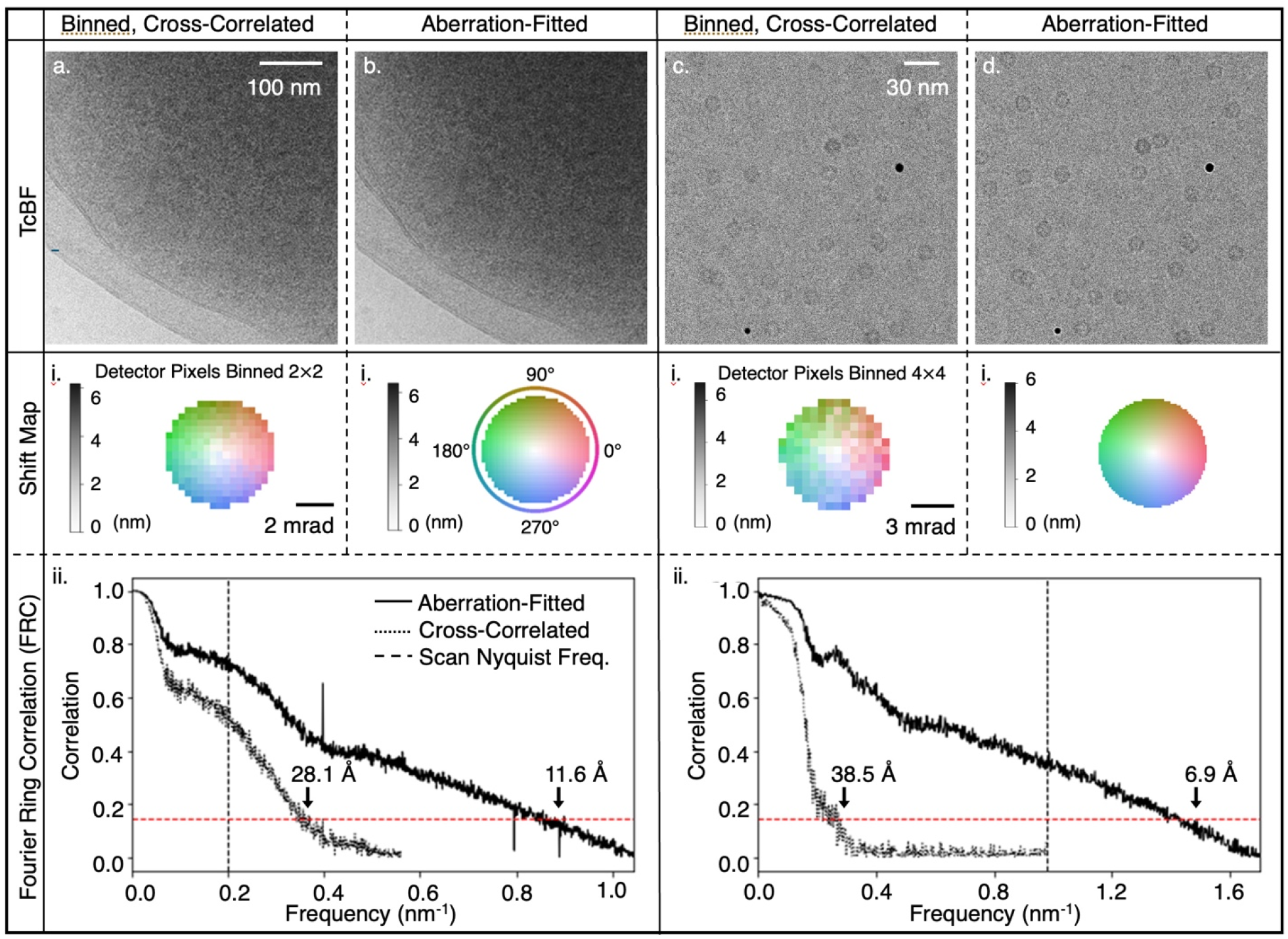
To overcome the low-SNR challenge for imaging frozen-hydrated apoferritin, 4-by-4 detector pixels are combined for successful cross-correlation (a). Fitting the shifts to the aberration function and applying the results to the original detector pixels help restore the information from individual detector pixels and improve image shift accuracy (b). The maps in the insets (i) present the shifts of images formed by each detector pixel, with the intensities indicating the magnitudes and the colors corresponding to the directions. The Fourier Ring Correlating (FRC) in the inset (ii) confirms the resolution enhancement by leveraging aberration, improving the cut-off resolution from 38.5 Å to 6.9 Å. For the thicker E.coli sample, the tcBF image with image shifts resolved on binned detector pixels (c) reveals the bilayer cell membrane and details in the interior of the cell. Leveraging aberration fitting results (d) pushes the resolution from 28.1 Å to 11.6 Å (inset (ii)). The 1/7 correlation threshold and the Nyquist sampling limit are labelled in the FRC plot. The images in (a) to (d) are cropped to show and the FRCs are computed with the full field of view. To calculate the FRC for tcBF-STEM, we generate two tcBF-STEM images from a single dataset by choosing alternating pixels within the BF disks and then reconstructing each subset independently. The normalized cross-correlation coefficient between the two resulting images represents the FRC of the dataset.

**Fig S10.**
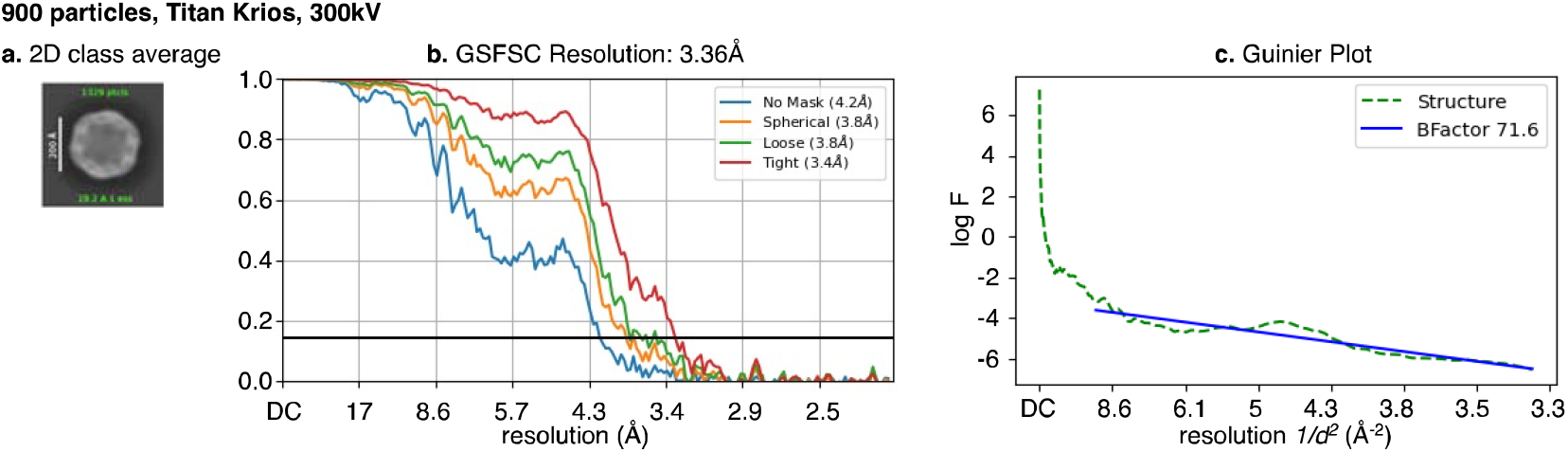
EFTEM ofVLP PP7 SPA with 900 particles reaches a nominal resolution of 3.36Å. The EFTEM data is acquired on a Cs-corrected TFS Krios at an accelerating voltage of300 kV. The image pixel size is 1.076 Å and the total dose is 52.18 e-/Å^2^ The defocus ranges from -lµm to -2µm. The final reconstruction is obtained with 900 particles and the analysis is done with cryoSPARC^44^

**Fig S11.**
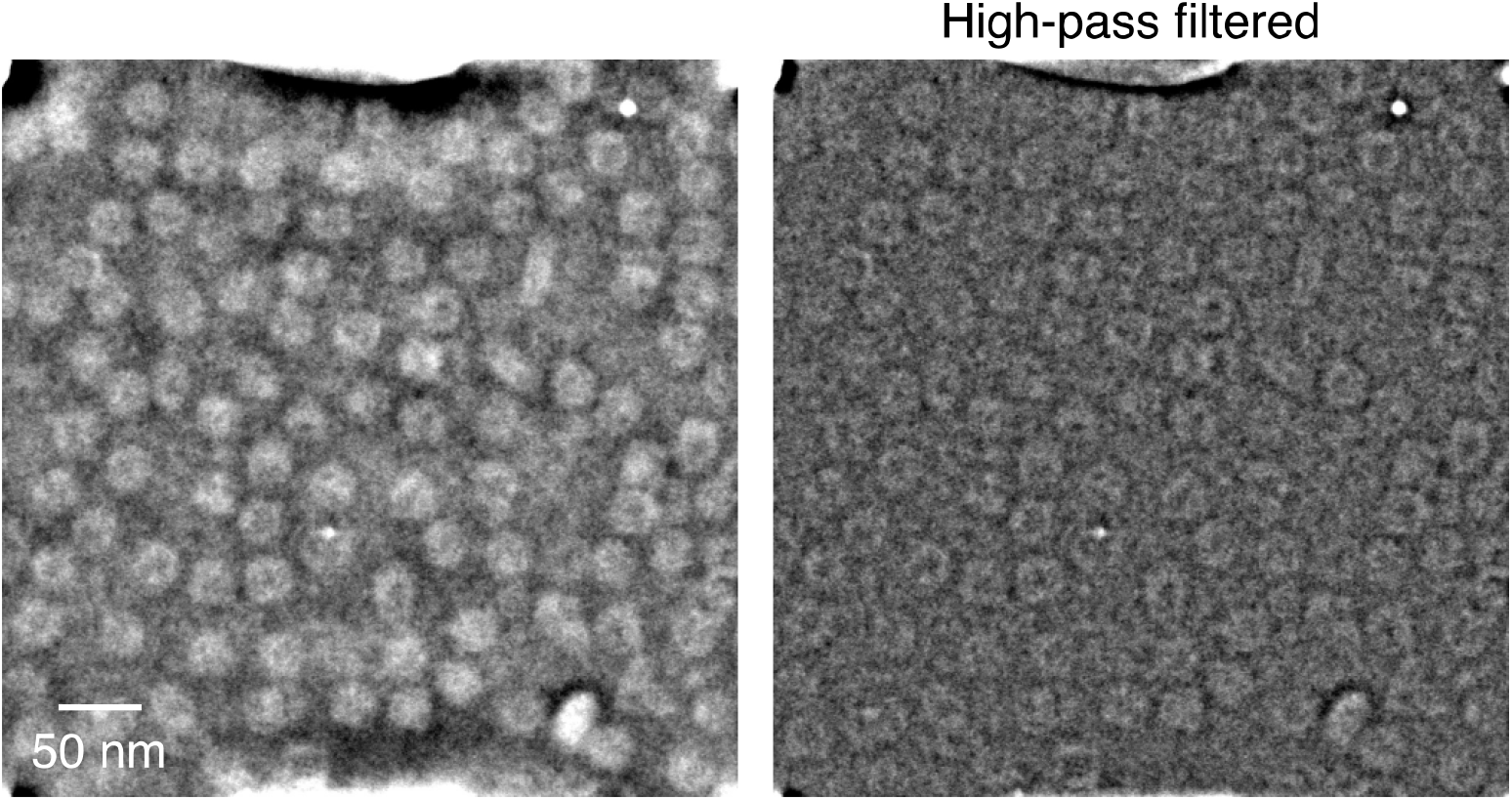
Cryo-iDPC on VLP PP7: the iDPC data was acquired using the TFS Panther detector on a TFS Spectra at 300 kV. Utilizing an in-house MATLAB code, iDPC images were generated (left), incorporating a high-pass filter (right) to eliminate low-frequency noise^51^.

